# Sharp cell-type boundaries emerge from temporal coordination between morphogen signals

**DOI:** 10.1101/2025.06.29.660399

**Authors:** Ruiqi Li, Yiqun Jiang, Sarah Platt, Tianchi Xin, Ryan Driskell, Kevin A. Peterson, Sarah Van, Hainan Lam, Shagun Lukkad, Eva-LaRue Barber, Chae Ho Lim, M. Mark Taketo, Yuval Kluger, Peggy Myung

## Abstract

Classic models of the French flag problem depict sharp cell-type boundaries emerging from threshold responses to morphogen gradients. Yet, how such boundaries arise during dynamic cell-state transitions remains unclear. We use hair follicle dermal condensates to study a sharp cell-type transition in which proliferative progenitors synchronously undergo cell-cycle exit and molecular differentiation. Using genetic and genomic approaches, we show that Wnt and Hedgehog signaling interact to coordinate the timing of these processes. When their activities are temporally aligned, intermediate transitional states are compressed through cell-cycle exit, producing a sharp cell-type boundary; when misaligned, transitional states expand, yielding fuzzy borders. Mechanistically, elevated Wnt activity promotes cell-cycle exit by regulating chromatin binding of the Hedgehog mediator GLI3. Hedgehog signaling induces differentiation genes in a Wnt-dependent manner and simultaneously elevates Wnt activity, aligning arrest and differentiation in time. These findings reveal a mechanism wherein morphogen interactions coordinate transitions to generate precise cell-type boundaries.

## Introduction

Developing tissues must position distinct cell types across space and time to assemble functional organs. How such patterns form was framed by Lewis Wolpert as the “French flag” problem, in which discrete cell types emerge at defined positions along a morphogen gradient (Fig. 1a)^1^. In this framework, graded morphogen signals are interpreted through concentration thresholds that delineate spatial domains of cell fate^2–6^. Yet, a fundamental question remains unresolved: how sharp cell-type boundaries emerge *in vivo* when the underlying signals and cell states vary continuously across a tissue. In developing systems, boundaries do not arise from instantaneous fate switches. Instead, cells progress through dynamic transitions and pass through intermediate states before committing to a new identity. Moreover, in many mammalian tissues, these fate transitions unfold while cells continue to proliferate, raising the question of how ongoing cell-cycle dynamics are coordinated with graded morphogen inputs. Thus, boundary sharpness may not only reflect where fate thresholds are crossed, but how rapidly cells traverse the same underlying developmental progression.

**Fig. 1:**
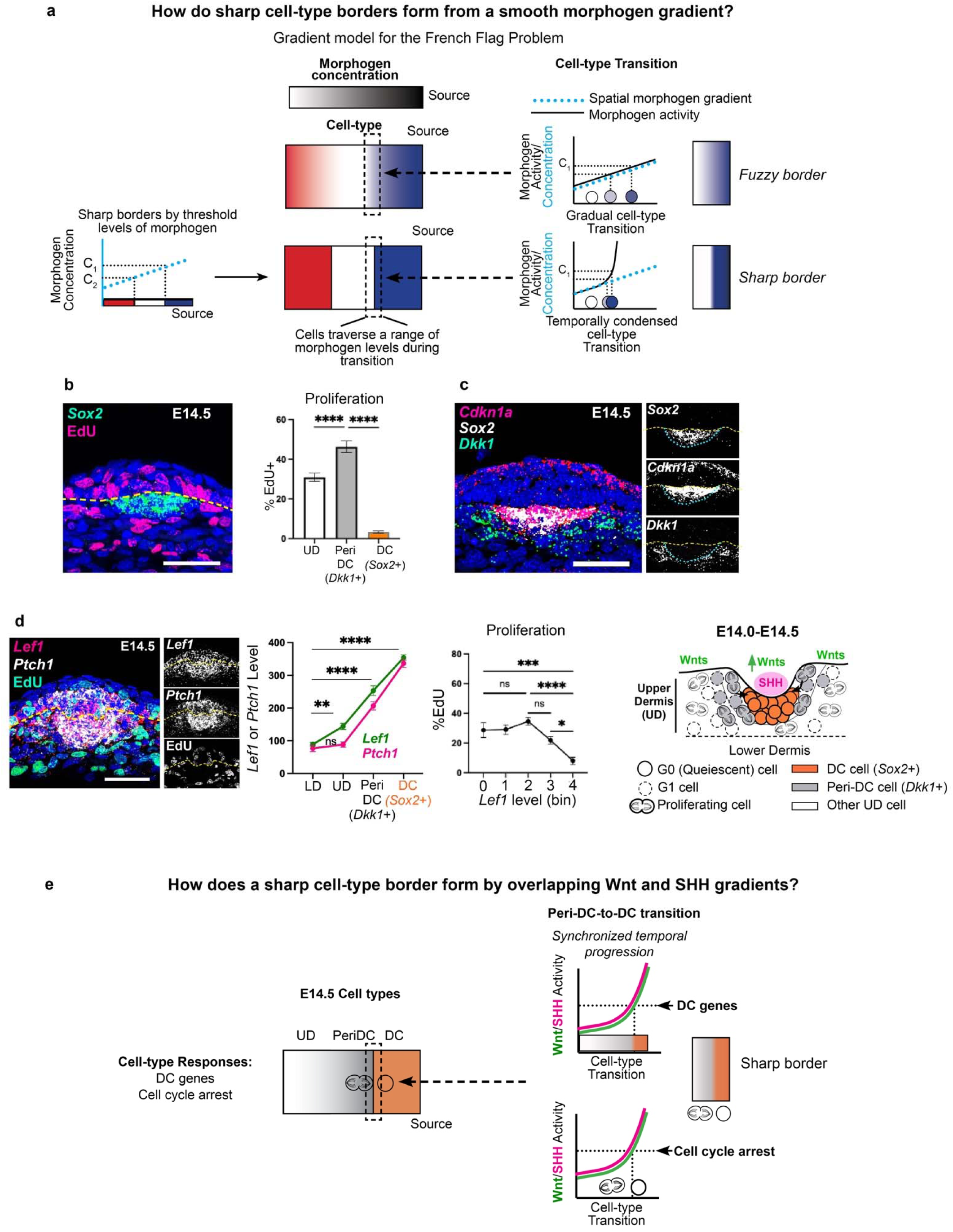
Cell-cycle exit and molecular differentiation are tightly coupled during dermal condensate formation. **a,** Left, illustration of the French flag problem with stripes representing distinct cell-type readouts positioned relative to a fixed morphogen source. Classical models propose that threshold concentrations of morphogen specify different cell-types with sharp borders. Right, schematic depiction of cellular responses to a linear morphogen gradient, illustrating either linear responses or non-linear transitions across morphogen levels. This contrasts static spatial threshold interpretations with a dynamic view in which cells traverse morphogen levels during fate transitions. **b,** FISH images of E14.5 mouse skin showing *Sox2*+ DCs form by E14.5, which lack EdU copositivity; quantification of %EdU by cell-type (UD, upper interfollicular dermis) (n=10). **c,** FISH image showing peri-DC (*Dkk1*+) cells adjacent to *Sox2*+ with *Cdkn1a* enriched at the DC boundary; (n=10). **d,** E14.5 skin stained for *Lef1* and *Ptch1* with transcripts quantified by spatial regions (LD, lower dermis) or %EdU+ by *Lef1* bin (n=5). **e,** Schematic summarizing dermal cell states with respect to spatial position and relative Wnt and SHH activity levels. DCs arise within regions of overlapping Wnt and SHH activity, where cell-cycle exit and DC gene expression occur within a narrow temporal window. Data as mean± SEM; \**P*<0.05, \*\**P*<0.01, \*\*\**P*<0.001, \*\*\*\**P*<0.0001, one-way ANOVA; ns, not significant. Scale bars=50 µm.

Addressing the temporal dimension of the French flag problem has been challenging, as most approaches to studying cell fate transitions emphasize cell origins or endpoints, rather than intermediate regulatory programs, or can visualize intermediate cell behaviors without molecular resolution^7–11^. We address this problem in a mammalian system by examining hair follicle dermal condensate formation, a uniquely tractable developmental transition in which the principal morphogen signals are known and cell-state progression occurs over a narrow temporal window^12–14^.

Dermal condensates are spheroid clusters of quiescent dermal cells that form beneath epithelial hair follicle placodes and are essential for hair follicle development^15–21^. At E13.5, prior to DC formation, Wnt signaling is broadly activated across the upper dermis, forming a gradient that is highest in cells closest to the epidermis^16, 22, 23^. Wnt activity is required but not sufficient for DC formation. Upon placode formation, epithelial cells secrete Sonic Hedgehog (SHH) and additional Wnt ligands, resulting in localized co-activation of Wnt and SHH in underlying dermal cells^21, 24–26^. Together, these signals are necessary and sufficient for DC formation, yet how Wnt and SHH signaling interact to drive the pre-DC-to-DC transition remains unknown, and DCs appear to emerge abruptly with few detectable intermediates.^8, 27, 28^.

Previous work has examined how tissue growth and proliferation affect morphogen gradients by shaping field size, ligand distribution, or the scaling of positional information^29–33^. In these frameworks, proliferation is typically considered in terms of how it modifies the signaling environment cells interpret. Many mammalian tissues, however, undergo fate specification while cells continue to divide, introducing an additional constraint, as proliferative expansion of intermediate states becomes a key factor influencing the precision of pattern formation. In this study, we focus on how interacting morphogen signals coordinate cell-cycle exit with differentiation to ensure spatially precise patterning in such proliferative contexts.

Here, we ask how interacting morphogen signals regulate the progression of a rapid cell-state transition to generate a sharp tissue boundary. By combining *in vivo* genetic perturbations, chromatin profiling, and a single-cell computational framework that disentangles concurrent biological programs, we are able to capture intermediate transcriptional states that are normally difficult to detect and to determine how their timing and spatial extent are regulated by these two pathways *in vivo*^34^. We show that Wnt and SHH differentially control cell-cycle exit and molecular differentiation, and that the temporal coordination of these processes determines whether transitional states are compressed or expanded, thereby controlling boundary sharpness. Mechanistically, we identify cross-regulation of transcription factor occupancy as a key mechanism linking signaling dynamics to cell-cycle exit.

## Results

### Cell-cycle exit and molecular differentiation are temporally coordinated during dermal condensate formation

Prevailing models of the French flag problem posit that distinct cell states are specified by threshold levels of morphogen concentration that vary with distance from a polarized source, predicting that sharp cell-type boundaries arise from spatial differences in morphogen signaling (Fig. 1a). To examine how this model applies to hair follicle dermal condensate (DC) formation, we asked how dermal cell types are spatially organized before and during DC formation. Prior to DC formation, dermal cells are proliferative and lack DC-specific markers (Extended Data Fig. 1a). Upon DC formation, DC cells express characteristic differentiation markers, including *Sox2*, *Sox18*, and *Gal*, and undergo cell-cycle exit, as indicated by minimal EdU nucleotide incorporation, while neighboring upper dermal cells that do not express DC genes remain highly proliferative. (Fig. 1b,c and Extended Data Fig. 1b,c).

Using *in vivo* labeling, we previously showed that DC cells arise from highly proliferative progenitors located in the upper dermis at E13.5 (Extended Data Fig. 1d)^16, 27^. As DC formation initiates, proliferating cells that give rise to DCs become localized to a region surrounding the nascent condensate (peri-DC region) where they express *Dkk1* (Fig. 1c and Extended Data Fig.1c). During the pre-DC–to–DC transition, *Dkk1*⁺ peri-DC cells exit the cell cycle while upregulating DC differentiation genes^27^. Consistent with this, DC marker expression overlapped with the *in vivo* G1/G0 Fucci2 reporter and with expression of the cell-cycle inhibitor *Cdkn1a* (*p21*), with only rare cells expressed *Cdkn1a* in the absence of *Sox2* (Fig. 1c and Extended Data Fig. 1e)^16, 35^. These data indicate that cell-cycle exit and DC differentiation occur concurrently during DC formation.

Given this temporal coincidence, we next asked how these two processes relate to Wnt and Sonic Hedgehog (SHH) signaling during DC formation. At E13.5, prior to SHH expression, Wnt signaling is active in the upper dermis (Extended Data Fig. 1a,b) but is not sufficient to induce DC gene expression^16, 22, 36^. Using *Lef1* as a canonical readout of Wnt activity, which is concordant with expression of multiple Wnt target genes (*Axin2*, *Wif1*, *Tcf7*) (Extended Data Fig. 1f)^16, 27^, we examined the relationship between Wnt signaling and proliferation. At E13.5, *Lef1* expression was highest in the most superficial layer of upper dermis and decreased across deeper dermal cell layers, yet this spatial pattern did not correlate with EdU incorporation, nor with DC gene expression (Extended Data Fig. 1a).

By E14.5, SHH expression in placode cells resulted in SHH activation in the underlying dermis. At this stage, dermal SHH activity (assessed by *Ptch1*) covaried with Wnt signaling (*Lef1*) across dermal populations, with significantly higher *Lef1* levels detected compared to E13.5 (Fig. 1d). The highest levels of both signals were observed in DC cells, intermediate levels in peri-DC (*Dkk1*⁺) progenitors, and progressively lower levels in surrounding upper dermal cells. In contrast to E13.5, elevated Wnt activity at E14.5 correlated with reduced proliferation, with the lowest EdU incorporation observed in DC cells (Fig. 1d,e). Thus, quiescence and molecular differentiation coincide spatially and temporally within a narrow domain characterized by elevated Wnt and SHH signaling.

### High Wnt activity induces cell-cycle exit independently of SHH signaling

To dissect the respective contributions of Wnt and SHH signaling to the pre-DC–to-DC transition, we perturbed each pathway prior to DC formation. Because both pathways are required for DC differentiation, standard loss-of-function approaches preclude analysis of their individual roles at this stage. Given that DC cells ultimately exhibit the highest levels of both Wnt and SHH activity, we asked whether these signals contribute to distinct components of the transition or act through a shared mechanism. We therefore examined the effects of elevating Wnt signaling at E13.5, prior to DC formation by expressing a constitutively active form of β-catenin (*Axin2CreER;βcat^f^*^l/EX3^, hereafter Actβcat; Fig. 2a, tamoxifen at E11.5).

**Fig. 2:**
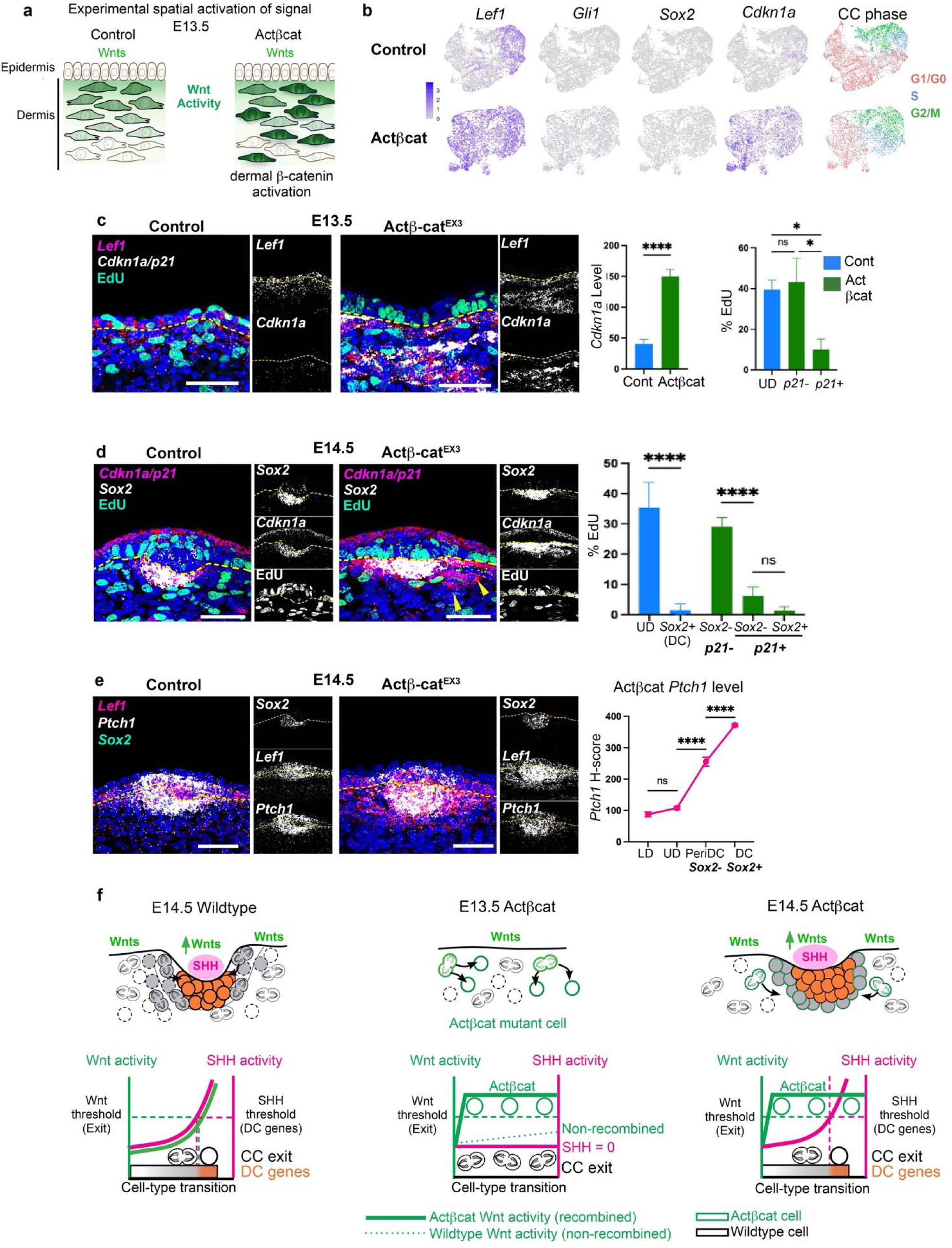
High Wnt activity induces cell-cycle exit independently of SHH signaling. **a,** Experimental setup in which β−catenin was constitutively activated in the dermis prior to DC formation (tamoxifen E11.5). **b,** UMAPs colored by indicated genes or cell cycle (CC) phase for E13.5 Actβcat mutant or paired control dermal scRNA-seq data (n=2). **c,** FISH showing E13.5 *Lef1*, *Cdkn1a*, EdU (left) and quantification of UD *Cdkn1a* transcripts (H-score) or %EdU of *Cdkn1a* (*p21*)+ or *p21*-cells in Actβcat UD vs. control UD (*p21*-) (n=3). **d,** FISH showing E14.5 *Sox2*, *Cdkn1a*, EdU (left) and quantification of %EdU+ cells in control or Actβcat *Cdkn1a*+ cells (almost all *Cdkn1a*+ cells co-express *Sox2* in control, n=5). **e,** E14.5 *Lef1*, *Ptch1*, *Sox2* FISH-stained sections (left) and quantification of *Ptch1* levels in Actβcat by cell type. **f,** Illustration showing that activated βcatenin (elevated Wnt activity) induces cell-cycle exit at E13.5 in the absence of SHH. At later stages, DC gene expression is restricted to cells experiencing high SHH activity with high Wnt activity. Data as mean± SEM; \**P*<0.05, \*\**P*<0.01, \*\*\**P*<0.001, \*\*\*\**P*<0.0001, one-way ANOVA; ns, not significant. Scale bars=50 µm.

To unbiasedly assess how increased Wnt activity reshapes dermal transcriptional states before DC differentiation, we performed single-cell RNA sequencing on E13.5 Actβcat and control dermis. Dermal cells were filtered for analysis using established markers (e.g., *Col1a1*, *PDGFRα*)^16, 27^. Consistent with prior work, Actβcat mutant dermal cells showed increased expression of Wnt target genes, including *Lef1*, *Axin2* and *Tcf7*, but not DC markers (e.g., *Sox2*) or SHH target genes (e.g., *Gli1*) (Fig. 2b). Notably, differential gene expression analysis revealed that *Cdkn1a*, a cell-cycle inhibitor normally expressed in DC cells at E14.5, was among the most significantly upregulated genes in E13.5 Actβcat dermal cells (Fig. 2b and Supplementary Table 1). Quantitative FISH confirmed elevated *Lef1* expression in E13.5 Actβcat dermal cells relative to controls and showed that Wnt target gene levels did not exceed those observed in wildtype DC cells, indicating that activated β-catenin elevates Wnt signaling but not above physiological levels (Fig. 2c and Extended Data Fig. 2a). FISH analysis further showed that constitutive Wnt activation was sufficient to induce expression of *Cdkn1a* (*p21*) at E13.5 in the absence of SHH activation, and *Cdkn1a*+ cells exhibited reduced EdU incorporation (Fig. 2c).

By E14.5, DCs were present in both control and Actβcat embryos but were enlarged in mutants (Extended Data Fig. 2b). In control embryos, *Cdkn1a* expression was largely restricted to *Sox2*+ DC cells, whereas in Actβcat mutants, it extended into surrounding *Sox2*-upper dermal cells (Fig. 2d). Both *Sox2*+ and *Sox2*− cells expressing high *Cdkn1a* exhibited low EdU incorporation (Fig. 2d). *Sox2*+ cells in Actβcat mutants were closest to placodes and expressed the highest levels of *Ptch1*, whereas surrounding mutant *Cdkn1a*+*Sox2*-cells expressed lower *Ptch1* levels, indicating that DC gene expression requires SHH signaling above a defined threshold (Fig. 2e). Thus, elevated Wnt signaling is sufficient to induce cell-cycle exit in the absence of SHH, whereas molecular DC differentiation requires additional SHH-dependent input (Fig. 2f).

### SHH signaling cell-autonomously elevates Wnt activity coincident with cell-cycle exit

Our results show that elevated Wnt activity induces cell-cycle exit without DC differentiation. We next examined how SHH signaling influences this transition. Specifically, we asked whether elevated SHH activity could initiate DC gene expression independently of cell-cycle exit. To address this, we expressed a constitutively activated form of Smoothened (*Axin2CreER;RosaLSL-Smom2YFP*, hereafter ActSMO) in the dermis prior to DC formation when Wnt signaling levels are low (tamoxifen E11.5; Fig. 3a). Whole-mount analysis of E14.5 skin showed that high SHH activation in Wnt-active dermal cells induced large dermal clusters expressing DC genes in the upper dermis (Fig. 3b). These clusters formed autonomously without placodes, indicating that DC gene expression can occur in the absence of placode-derived Wnt ligands^27^. Notably, many cells in the outer layers of Sox2+ dermal clusters remained proliferative, as indicated by %Sox2*+*/EdU+ double labeling, demonstrating that DC gene expression can occur independently of cell-cycle arrest. By contrast, Sox2+ cells in the center of clusters lacked EdU incorporation similar to quiescent DC cells in wildtype embryos.

**Fig. 3:**
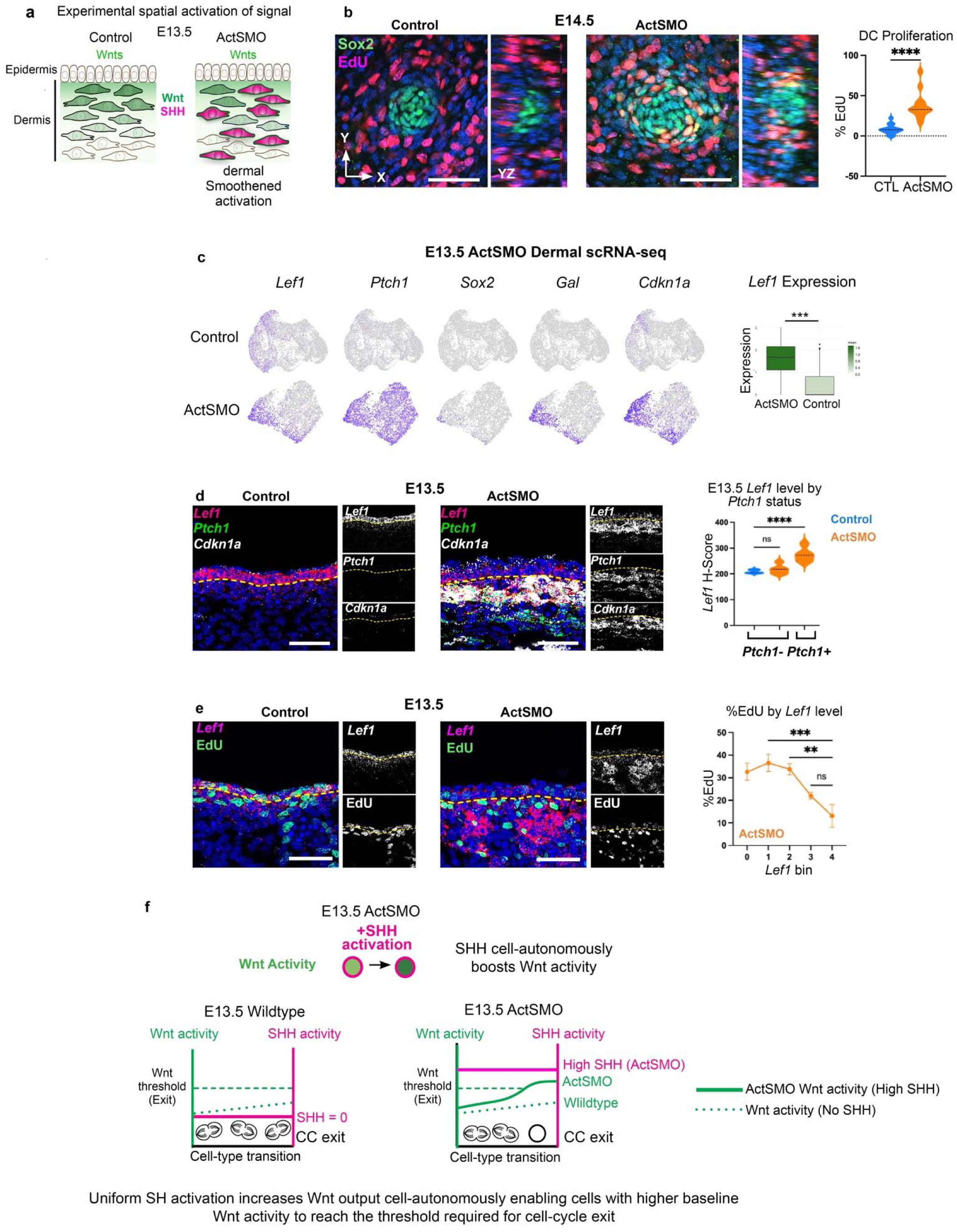
SHH signaling cell-autonomously elevates Wnt activity in association with cell-cycle exit. **a,** Experimental setup showing expression of constitutively activated Smoothened in dermal cells prior to DC formation when Wnt activity is low (tamoxifen E11.5). **b,** Whole-mount images of E14.5 ActSMO and control skin stained with EdU and Sox2 with quantification of %EdU+ of Sox2+ cells (n=8). **c,** UMAPs of E13.5 control and ActSMO dermal scRNA-seq data with indicated gene expression; right, quantification of scRNA-seq *Lef1* levels in UD cells. **d,** E13.5 control or ActSMO mutant skin stained for *Lef1*, *Ptch1* and *Cdkn1a* with quantification of average *Lef1* expression by *Ptch1* expression status (n=8). **e,** E13.5 skin showing ActSMO boosts *Lef1* gene expression which corresponds with decreased EdU incorporation (%EdU). **f,** Cartoon illustrating that high SHH activity cell-autonomously elevates Wnt signaling, and elevated Wnt activity coincides with cell-cycle exit. Data as mean± SEM; \**P*<0.05, \*\**P*<0.01, \*\*\**P*<0.001, \*\*\*\**P*<0.0001, one-way ANOVA. Scale bars, 50 µm.

Given the observed quiescent central Sox2+ dermal cells (Fig. 3b), we asked whether there were similar molecular changes to those seen in the E13.5 Actβcat mutant. We obtained scRNA-seq data from E13.5 ActSMO and paired control skin (Fig. 3c). As expected, E13.5 ActSMO dermal cells exhibited uniformly high *Ptch1* expression. *Lef1* expression remained largely enriched in upper dermal (UD) cells but was significantly elevated compared to E13.5 controls (Fig. 3c,d). FISH analysis confirmed increased *Lef1* expression in E13.5 ActSMO dermis despite the absence of placodes (Fig. 3d). Importantly, only *Ptch1*+ cells (recombined) showed elevated *Lef1* expression, while neighboring *Ptch1*-negative cells (non-recombined) retained normal *Lef1* levels comparable to E13.5 control UD cells, indicating a cell-autonomous effect. ActSMO cells with high *Lef1* expression also exhibited elevated *Cdkn1a* expression and low rates of EdU incorporation (Fig. 3d,e). Together, these findings show that SHH activation elevates Wnt signaling cell-autonomously and is associated with induction of cell-cycle exit, independent of placode Wnt ligands (Fig. 3f).

### GeneTrajectory resolves transcriptional programs associated with cell-cycle exit

In the ActSMO condition, DC genes are expressed in cells that remain proliferative (Fig. 3b), indicating that cell-cycle exit and DC gene expression can be temporally uncoupled. To resolve the gene programs underlying these processes, we applied GeneTrajectory (GT), a computational approach that separates co-occurring transcriptional programs from whole transcriptome scRNA-seq data^34^. Unlike conventional pseudotime methods that assign cells to a single lineage trajectory, GT directly constructs gene trajectories, allowing individual cells to be analyzed across multiple overlapping biological processes.

We first applied GT to E14.5 wildtype dermal scRNA-seq data, identifying three gene trajectories, corresponding to proliferation (S/G2/M), DC differentiation and lower dermal differentiation (Fig. 4a). The DC trajectory was partitioned into stages using gene bins, and cells were visualized by gene bin scores (Fig. 4a and Extended Data Fig. 3a)^34^. *Lef1* and *Ptch1* expression covaried across DC trajectory stages, with maximal expression at the terminal stage 6 (Fig. 4b and Extended Data Fig. 3b). Canonical DC genes, including *Gal*, *Sox18* and *Sox2,* were similarly enriched at stage 6 (Fig. 4c). Cell-cycle analysis revealed that stage 6 cells were quiescent with a G1 fraction near 1.0, and elevated *Cdkn1a* expression (Fig. 4c). Thus, in the wildtype condition, cell-cycle exit and molecular DC differentiation coincide at the terminus of the DC trajectory.

**Fig. 4:**
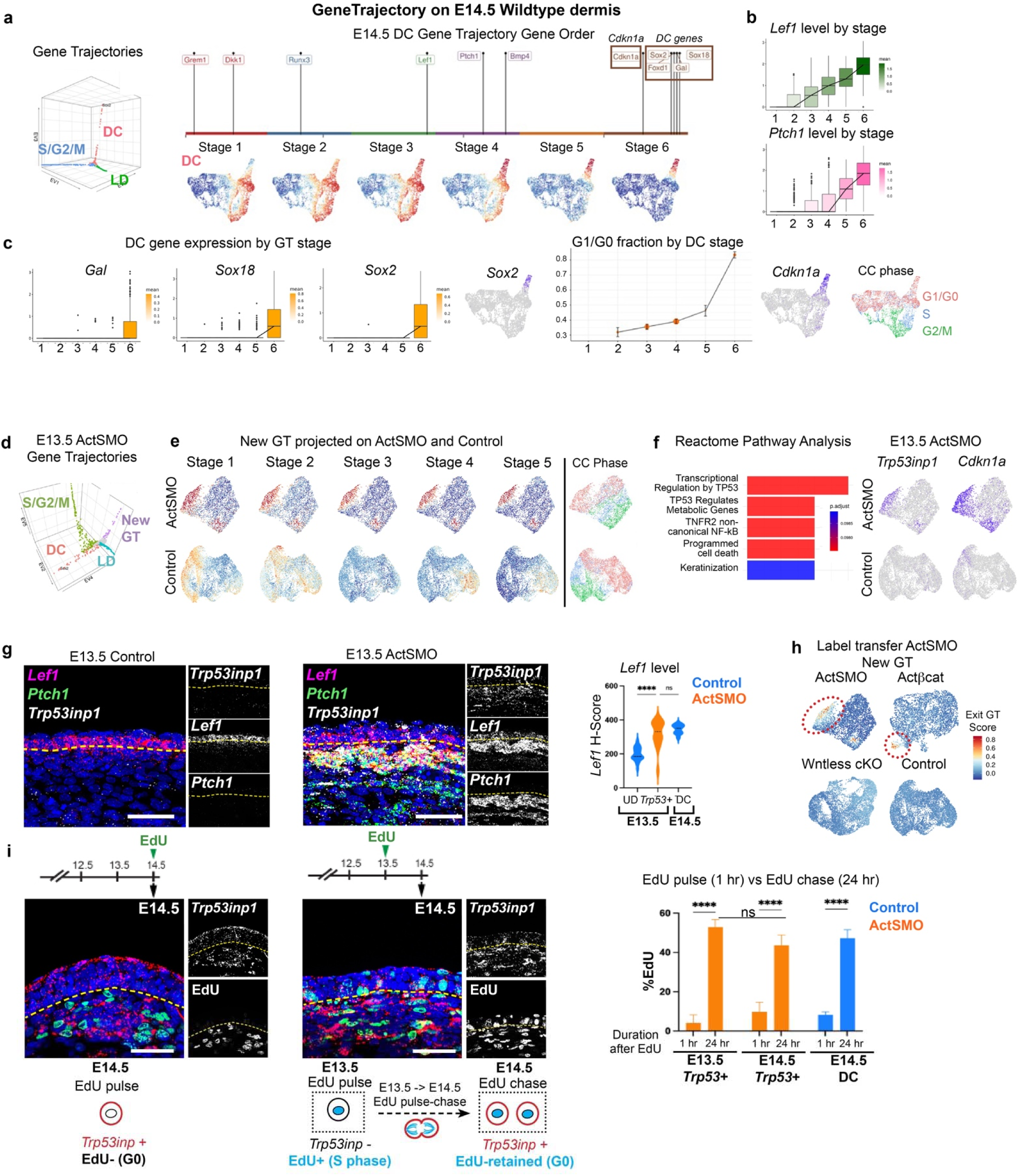
GeneTrajectory defines transcriptional changes accompanying cell-cycle exit. **a,** GeneTrajectory diffusion map and gene order of E14.5 wildtype DC GT with dermal UMAPs colored by DC GT stages. **b,** *Lef1* and *Ptch1* levels by DC GT stage. **c,** DC gene expression by DC GT stage. Right, G1/G0 fraction by DC GT stage with UMAP of *Cdkn1a* expression or CC phase. **d,** E13.5 ActSMO GT diffusion map. **e,** ActSMO and control dermal UMAPs colored by expression of the mutant New GT stages or CC phase. **f,** Pathway analysis of genes from new mutant GT and UMAPs of control and mutant dermal cells colored by indicated genes. **g,** FISH images of indicated genes in E13.5 control or ActSMO skin with quantification of *Lef1* expression in control E13.5 upper dermal (UD) or E14.5 DC cells vs. E13.5 ActSMO *Trp53inp1*+ cells (n=3). **h,** Label transfer of the E13.5 ActSMO new GT onto indicated E13.5 mutant or control dermal populations showing cells that express this similar gene module. **i,** EdU pulse-chase experiments of E13.5-E14.5 ActSMO mutant and corresponding quantification of %EdU of *Trp53inp1*+ cells (n=3). Bottom cartoon illustrating *Trp53inp1*+ cells undergo cell cycle exit as immediately after a division. Data as mean ± SEM, one-way ANOVA. Scale bars, 50 µm.

We next applied GT to E13.5 ActSMO scRNA-seq data in which cell-cycle exit and DC differentiation are uncoupled. GT identified four trajectories (Fig. 4d and Extended Data Fig. 3c). Two were similar to the control, corresponding to proliferation (S/G2/M) and LD differentiation. Two additional trajectories were unique to the ActSMO dermis: one containing DC genes and a second lacking DC markers. Cells expressing this latter program localized to a subset of Wnt-active cells and were largely restricted to G1/G0 phase (Fig. 4e and 3c). Pathway analysis revealed enrichment for TP53-associated cell-cycle inhibitory genes, including *Trp53inp1* and *Plk2,* overlapping with *Cdkn1a* expression (Fig. 4f and Supplementary Table 1). FISH analysis confirmed significant upregulation of both *Cdkn1a* and *Trp53inp1* in the E13.5 ActSMO dermis relative to controls (Fig. 4g and 3d and Extended Data Fig. 3d)

In ActSMO mutants, SHH-active cells expressing *Trp53inp1* exhibited significantly higher *Lef1* levels than control upper dermal cells (Fig. 4g), consistent with earlier results showing that SHH elevates Wnt activity and promotes quiescence (Figs. 3d and 2c). To test whether elevatd Wnt signaling alone induces this TP53-associated program, we applied GT to the E13.5 Actβcat dermis, which induces *Cdkn1a* but not DC genes (Extended Data Fig. 3e). GT analysis identified a similar *Trp53inp1*-containing program in E13.5 Actβcat mutants (Supplementary Table 1). We applied a module projection (label transfer) method to assess whether the gene program was shared across different perturbations. Specifically, we transferred the E13.5 ActSMO-specific trajectory onto E13.5 Actβcat and control datasets, which revealed a shared molecular state in both mutants that was absent in controls (Fig. 4h).

To determine whether this program reflects a general consequence of reduced proliferation, we analyzed E13.5 Wntless conditional knockout (*K14Cre;Wls^fl/fl^*) embryos in which epidermal Wnt ligand secretion is disrupted, resulting in loss of dermal Wnt signaling and reduced dermal proliferation (Extended Data Fig. 3f)^22, 34, 37^. Label transfer of the ActSMO/Actβcat-specific trajectory was not detected in Wntless cKO dermis (Fig. 4h and Extended Data Fig. 3f), indicating that this TP53-associated program is Wnt-dependent and specific to DC-associated cell-cycle exit.

Finally, we examined the cell-cycle dynamics of cells expressing this gene program. At E13.5 and E14.5, most ActSMO *Trp53inp1*+ cells showed low EdU incorporation comparable to quiescent DC cells (Fig. 4i and Extended Data Fig. 4a). EdU pulse-chase experiments demonstrated that a substantial fraction of *Trp53inp1*+ cells retained EdU nucleotide similarly to wildtype DCs (Fig. 4i and Extended Data Fig.1d)^16, 27^. Although ActSMO dermal clusters contained proliferating Sox2+ cells at E14.5, most became quiescent by E15.5 (Extended Data Fig. 4b), indicating that these proliferating cells exit the cell cycle after division. Together, these data define a shared transcriptional program underlying cell-cycle exit during the pre-DC-to-DC transition, which we term the *“Exit” gene trajectory*.

### Loss of GLI3 chromatin binding induces cell-cycle exit downstream of elevated Wnt signaling

Although Wnt activity is required for dermal proliferation, our results indicate that elevated Wnt activity promotes cell-cycle exit during DC formation^16, 22^. To examine the underlying mechanism, we profiled genes differentially bound by CTNNB1 (β−catenin) in E13.5 Actβcat and control dermis. CTNNB1 binding was enriched at canonical Wnt target genes (e.g., *Sp5*, *Axin2*, *Lef1*), but not at *Cdkn1a,* Exit GT genes, or other loci associated with cell cycle inhibition (Extended Data Fig. 5a,b)^22^. These findings raised the question of how elevated Wnt activity promotes cell-cycle exit despite the absence of direct CTNNB1 binding at cell-cycle inhibitory genes. Given that Wnt activity is elevated upon SHH activation, it was plausible that components of the SHH pathway mediate Wnt-dependent cell-cycle exit. In other tissues, Wnt signaling regulates *Gli3* expression independently of SHH^38^. In the absence of SHH ligand, Gli3 is predominantly present in its cleaved repressor form, which binds a broad range of SHH target genes^39, 40^. FISH analysis showed that *Gli3* was expressed in Wnt-active dermal cells in both control and Actβcat embryos at E13.5 prior to SHH expression (Fig. 5a). To profile GLI3 binding, we performed CUT&RUN in E13.5 Actβcat and control dermis using *Gli3Flag* knockin embryos. Interestingly, GLI3 occupancy was globally reduced in the Actβcat condition as indicated by decreased GLI3-bound reads at transcriptional start sites (Fig. 5b). De novo motif analysis confirmed enrichment of GLI motifs at regions showing reduced binding (Extended Data Fig. 5c)^41^. These regions were enriched for genes associated with SHH signaling, despite the fact that most canonical SHH target genes (e.g., *Ptch1*, *Gli1*) were not expressed and lacked active histone marks (H3K4me3 or H3K27ac) at E13.5 (Fig. 5c). These observations are consistent with a recent report indicating that GLI3 binding prior to SHH does not actively repress transcription (i.e., is inert) and that many targets become activated only after SHH expression^42^. Western Blot analysis confirmed that cleaved Gli3 protein levels were retained in the Actβcat mutant, showing that reduced chromatin occupancy was not due to loss of Gli3 protein and that Gli3 remained predominantly in its cleaved repressor form (Fig. 5d). These results show that elevated Wnt signaling alone reduces GLI3 chromatin occupancy at many genomic regions independently of SHH.

**Fig. 5:**
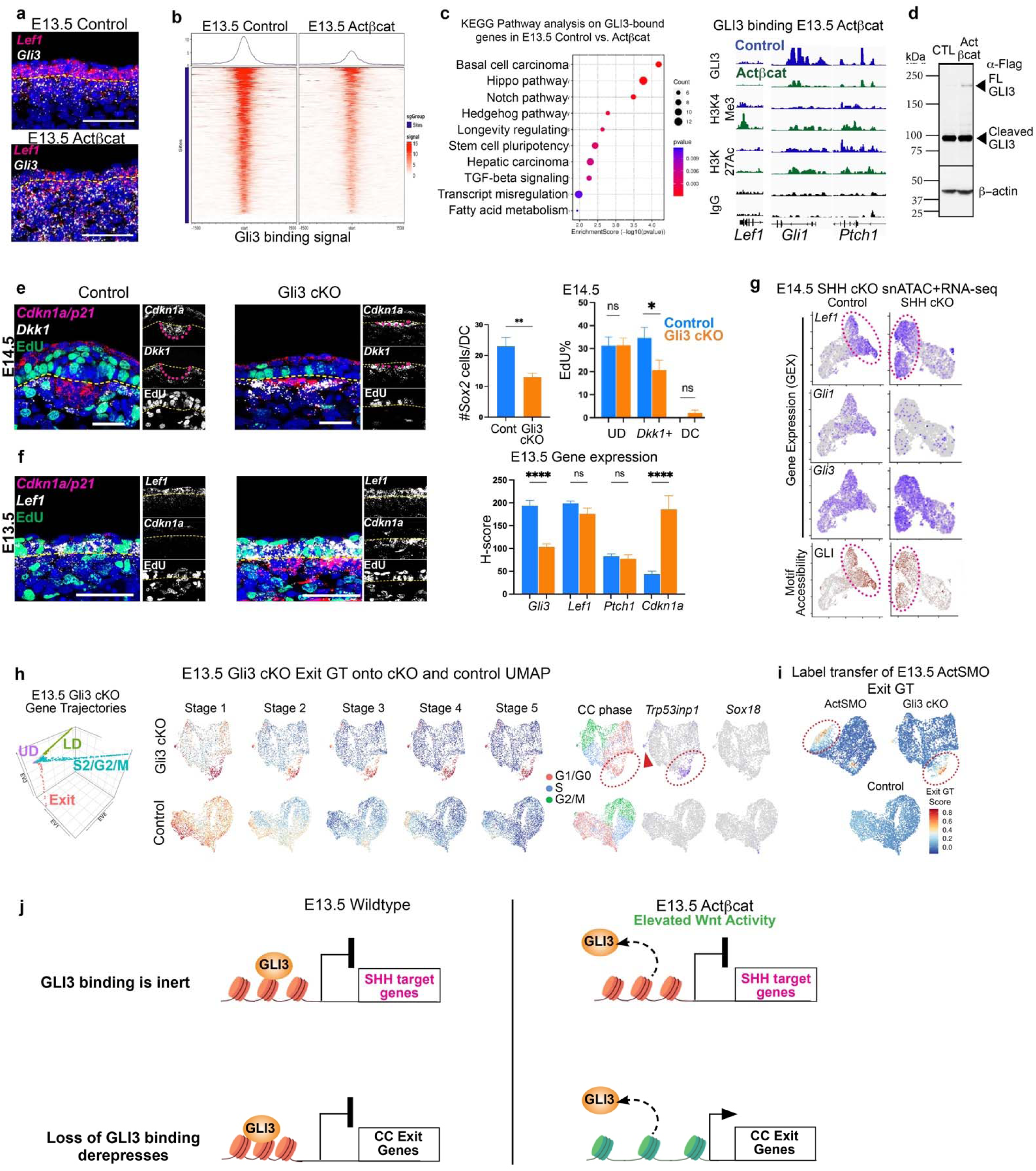
Loss of GLI3 chromatin binding induces premature cell-cycle exit downstream of elevated Wnt signaling. **a,** *Gli3* expression in E13.5 control and Actβcat skin prior to SHH. **b,** Heat map of GLI3 binding at all transcriptional start sites (TSS) in E13.5 control or Actβcat skin. **c,** KEGG pathway analysis of GLI3 unbound regions in the Actβcat mutant. Right, Genome browser tracks showing binding signal of respective antibodies at indicated genes. **d,** Western blot of cell lysates from control or Actβcat mutant skin stained with anti-Flag (Gli3) or β-actin (n=3). **e,** FISH of E14.5 control and Gli3 dermal cKO (tamoxifen E11.5) stained for indicated genes; quantification of DC size and %EdU of control or Gli3 cKO by cell type (n=3). **f,** FISH of E13.5 control and Gli3 cKO stained for indicated genes with quantification of transcripts by FISH (n=4). **g,** snATAC+RNA-seq UMAPs of co-embedded E14.5 control and epidermal SHH cKO showing either gene expression or GLI motif accessibility. **h,** GT diffusion map of E13.5 Gli3 cKO dermal cells, showing an Exit GT. UMAPs of control or Gli3 cKO colored by Exit GT bins, indicated genes or cell-cycle phase. **i,** Label transfer of E13.5 ActSMO Exit GT onto Gli3 cKO or control scRNA-seq data showing a shared Exit gene GT. **j,** Elevated Wnt activity reduces GLI3 chromatin occupancy, derepressing genes that mediate cell-cycle exit, but not DC differentiation. Here, canonical SHH target genes and DC differentiation genes remain inactive in the absence of SHH signaling. Upon co-activation of Wnt and SHH, canonical SHH targets and DC genes are induced. Data as mean± SEM; \**P*<0.05, \*\**P*<0.01, \*\*\**P*<0.001, \*\*\*\**P*<0.0001, one-way ANOVA; ns, not significant; scale bars, 50 µm.

To test whether loss of GLI3 binding is sufficient to induce cell-cycle exit, we ablated *Gli3* in the dermis (*Axin2CreER;Gli3^fl/fl^* or Gli3 cKO; tamoxifen E11.5). At E14.5, DCs in Gli3 cKO embryos were smaller than in controls (Fig. 5e), and *Dkk1*+ peri-DC progenitors showed premature upregulation of *Cdkn1a* and reduced EdU incorporation (Fig. 5e and Extended Data Fig. 5d). Similarly, at E13.5 prior to DC formation, Gli3 cKO dermal cells exhibited elevated *Cdkn1a/p21* expression and reduced proliferation, phenocopying the Actβcat condition (Fig. 5f and Extended Data Fig. 5e).

FISH analysis confirmed reduced *Gli3* expression in the dermis of E13.5 Gli3 cKO embryos, while Wnt target genes were unaffected, suggesting that Gli3 acts downstream of elevated Wnt activity (Fig. 5f and Extended Data Fig. 5f). Canonical SHH target genes (e.g. *Ptch1*) were also not induced, confirming that loss of Gli3 does not broadly derepress SHH targets and that the lack of SHH target expression in the Actβcat condition is unlikely due to residual GLI3 repression. To assess whether GLI3 binding is associated with chromatin organization, we performed chromatin capture (Micro-C) in E13.5 wildtype dermis. GLI3-bound loci (e.g., *Ptch1*) were actually modestly enriched in A (open) rather than B (closed) compartments but rarely participated in chromatin loops (<1%) and lacked active promoter marks (Extended Data Fig. 5g). By contrast, canonical Wnt target genes (e.g., *Lef1* and *Sp5)* formed chromatin loop interactions and showed active histone modifications (Extended Data Fig. 5g). Moreover, snATAC+RNA-seq analysis of E14.5 embryos that lack epidermal *Shh* showed significant GLI motif accessibility localized to Wnt-active cells even without Gli activator function (Fig. 5g). Together, these data indicate that GLI3 does not actively repress many bound regions and may instead prime GLI sites for accessibility prior to SHH.

As expected, scRNA-seq analysis of E13.5 Gli3 cKO dermis showed reduced *Gli3* expression. Neither DC nor canonical SHH target genes were detected, and Wnt target genes were retained (Extended Data Fig. 6a,b). GT analysis identified four trajectories in Gli3 cKO dermis, three corresponding to control populations and a fourth absent in controls (Fig. 5h and Extended Data Fig.6c). This trajectory overlapped with the Exit GT identified in the Actβcat and ActSMO mutants and was enriched for cell-cycle inhibitory genes, including *Trp53inp1* (Fig. 5h and Supplementary Table 1). Additionally, label transfer of the Exit GT from the ActSMO mutant to the Gli3 cKO dataset identified a corresponding population that expressed this gene program (Fig. 5i), showing that this Exit gene program is shared across ActSMO, Actβcat, and Gli3 cKO conditions. Using EdU pulse-chase experiments, we found that *Cdkn1a*-high cells in the Gli3 cKO were largely quiescent, while representing recent progeny of cell divisions (Extended Data Fig. 5e). Thus, β-catenin and Gli3 function in a common pathway regulating cell-cycle exit.

Cross-referencing genes that lost GLI3 binding in E13.5 Actβcat dermis with those upregulated in E13.5 Gli3 cKO dermis identified multiple cell-cycle inhibitory genes, including *Mxd4* and *Efna5*, which showed increased expression and more active promoter marks (Extended Data Fig. 6d). In addition, Exit GT genes such as *Baiap2* and *Cpt1c* also exhibited reduced GLI3 binding and increased expression. Together, these data indicate that β-catenin and Gli3 function within a shared pathway regulating cell-cycle exit in which elevated Wnt signaling causes loss of GLI3 chromatin occupancy and induction of genes associated with cell-cycle exit (Fig. 5j).

### Uniformly high SHH activity induces dermal condensate gene expression across the Wnt gradient

High SHH activity can induce DC gene expression before cell-cycle exit, indicating that DC differentiation does not strictly require a preceding quiescent state (Fig. 3b). However, in the wildtype condition, DC genes are induced abruptly as cells exit the cell cycle, suggesting that Wnt activity contributes to the timing of DC gene induction. Given that SHH cell-autonomously elevates Wnt activity, we examined whether Wnt signaling levels modulate how DC genes are induced in response to high SHH activity.

To address this, we analyzed the DC gene trajectory in the E13.5 ActSMO mutant, in which SHH activity is uniformly high across cells spanning a range of Wnt activity (Fig. 6a and Extended Data Fig 7a). In the E14.5 wildtype DC gene trajectory, DC genes are induced primarily at the terminal stage, coinciding with the highest levels of *Lef1* and *Ptch1* expression (Fig. 4a,c). By contrast, in the ActSMO condition, DC genes, including *Gal*, *Sox18,* and *Sox2*, were induced progressively across DC GT stages, with cumulative expression correlating with increasing *Lef1* levels (Fig. 6b,c,d and Extended Data Fig. 7b). Cells with highest *Lef1* levels showed the most advanced progression along the trajectory and the strongest DC gene expression (Fig. 6c,d).

**Fig. 6:**
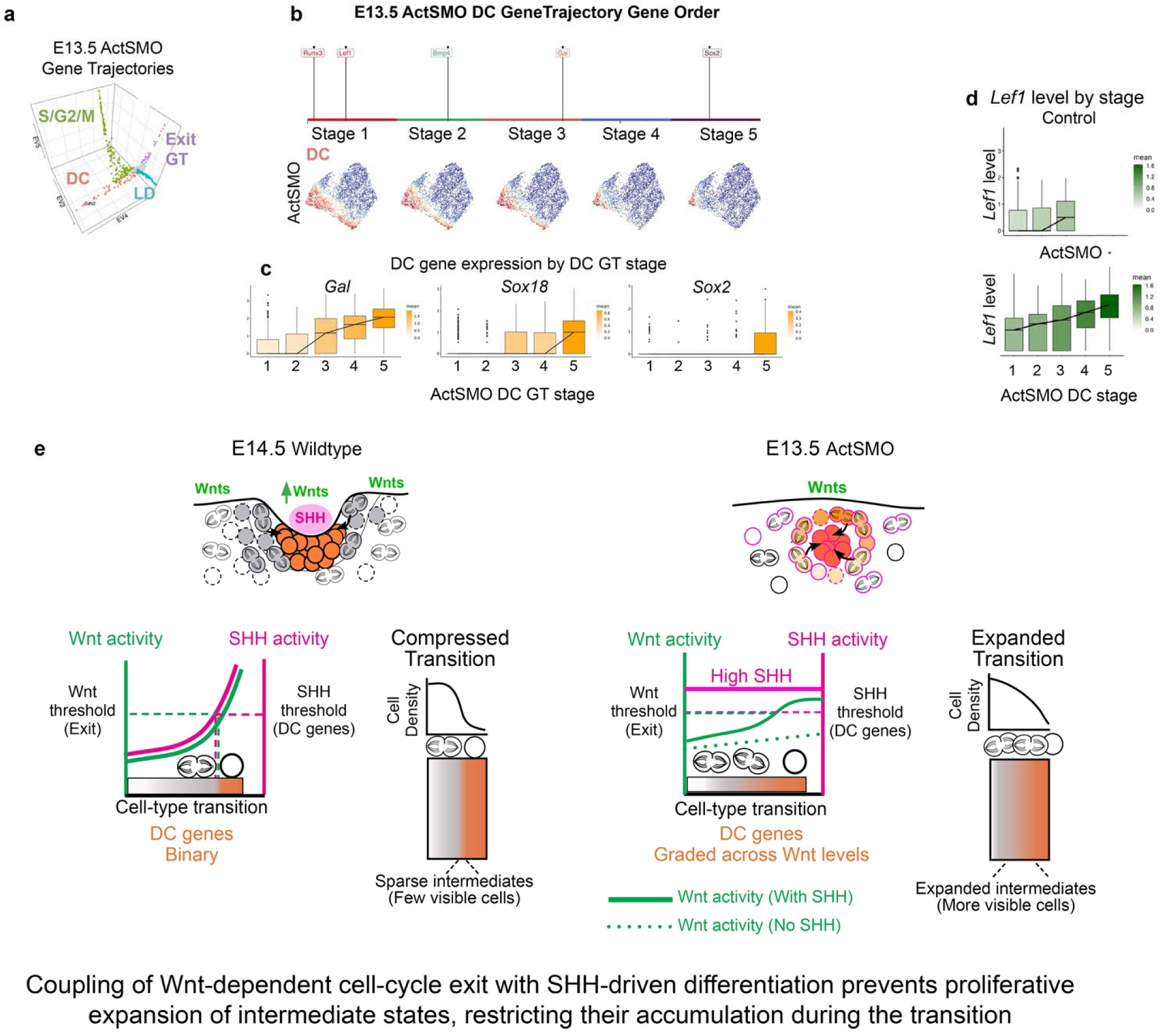
Uniformly high SHH activity induces dermal condensate gene expression across the Wnt gradient. **a,** E13.5 ActSMO dermal GT diffusion map. **b,** Gene order (top) of ActSMO DC GT, showing DC genes expressed across the trajectory; expression of DC GT stages colored onto ActSMO dermal UMAPs. **c,** Expression levels of DC genes by DC GT stage. **d,** *Lef1* levels by DC GT stage of ActSMO and control. **e,** Cartoon illustrating that uniform SHH activation permits DC gene expression across a range of Wnt activity levels, resulting in sequential induction of DC genes that correlates with *Lef1* levels. Here, DC genes can be expressed in proliferating cells of lower Wnt activity, expanding intermediate states before coordinated cell-cycle exit is achieved. The orange/gray bars represent the extent of intermediate DC transcriptional states along the trajectory and not a direct measure of temporal kinetics.

Consistent with the requirement for Wnt signaling in DC differentiation, cells with high SHH activity but lacking Wnt signaling in the ActSMO condition failed to express DC genes (Fig. 6c and Extended Data Fig. 7c)^27^. Thus, uniform SHH activation expands the window over which Wnt-competent cells initiate DC gene expression, resulting in graded rather than abrupt induction of DC genes. In both E14.5 wildtype and Actβcat (high Wnt) conditions, DC genes were restricted to Wnt-active cells with high SHH activity, indicating that DC differentiation requires SHH signaling above a defined threshold (Fig. 6e).

### Coupling Wnt and SHH gradients is sufficient to synchronize the timing of cell-cycle exit with molecular differentiation

Given that cell-cycle exit and DC differentiation coincide where high Wnt activity overlaps with SHH signaling, we tested whether coupling these gradients is sufficient to restore temporal coordination between the two processes. In wildtype skin, this overlap occurs in dermal cells adjacent to placodes. We therefore induced epidermal SHH expression at E13.5 (*K14Cre;Rosa-LSL-SHH*; SHH OE) to activate SHH signaling in Wnt-active dermal cells prior to placode formation (Fig. 7a)^43, 44^. FISH analysis revealed that *Lef1* and *Ptch1* covaried with dermal depth with the highest levels in cells closest to the epidermis (Fig. 7b). Compared to E13.5 controls, *Lef1* levels were increased in the superficial dermis, indicating that SHH ligand expression enhanced Wnt activity. DC genes (e.g., *Sox2*, *Gal*) were restricted to these superficial layers and coincided with *Cdkn1a* expression and low EdU incorporation (Fig. 7c and Extended Data Fig. 8a). *Dkk1* expression localized to a deeper layer beneath cells expressing DC genes. scRNA-seq analysis of E13.5 SHH OE dermis confirmed overlapping *Lef1* and *Ptch1* expression, with DC genes and *Cdkn1a* restricted to cells with the highest combined signal levels (Fig. 7d).

**Fig. 7:**
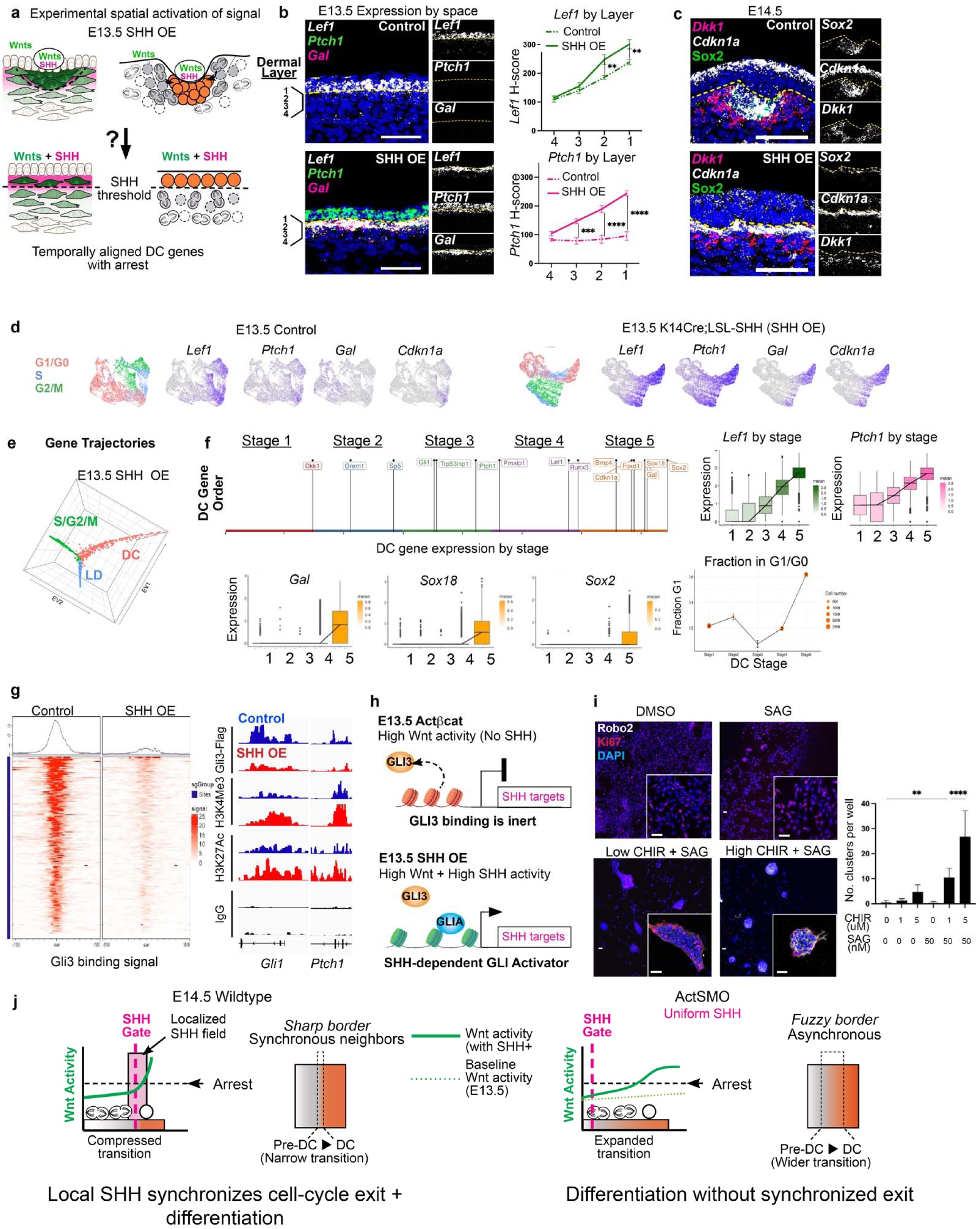
Coordinated Wnt and SHH signaling synchronizes cell-cycle exit and dermal condensate differentiation. **a,** Experimental design to test whether coordinated Wnt and SHH activity is sufficient to synchronize cell cycle exit with DC genes by co-expressing Wnt and SHH ligands across the epidermis at E13.5 (*K14Cre;Lox-STOP-Lox-Shh*; SHH OE). **b,** FISH images of E13.5 control and mutant showing covarying levels of *Ptch1* and *Lef1* across vertical dermal layers in the SHH OE mutant and elevated *Lef1* levels in layers where *Ptch1* is concurrently activated. **c,** E14.5 control and SHH OE with spatial partitioning of cells by cell type genes and *Cdkn1a* coinciding with *Sox2* expression (n=3). **d,** E13.5 control or SHH OE UMAPs with indicated genes or cell-cycle phase. **e,** GT diffusion map of E13.5 SHH OE dermal scRNA-seq data. **f,** DC GT gene order of SHH OE (top) with expression of DC genes and fraction G1/G0 by DC GT stage (bottom) showing coincident induction of DC genes and cell-cycle exit at the terminal stage. Right, covarying *Lef1* and *Ptch1* levels by DC GT stage. **g,** Heat map of GLI3 binding at TSS in E13.5 control versus SHH OE. Genome browser tracks showing reduced GLI3 binding and active histone marks at *Gli1* and *Ptch1*. **h,** Cartoon showing loss of GLI3 binding upon Wnt and SHH coactivation with activation of canonical SHH targets. **i,** Confocal images of 3D dermal cultures stained with Robo2 (DC marker) showing Robo2+ clusters lacking Ki67 in high CHIR+SAG condition. Right, dermal cluster number in 3D cultures by condition. **j,** Model illustrating how coordination between Wnt and SHH signaling regulates the timing of cell-cycle exit and molecular differentiation during the pre-DC-to-DC transition. When morphogen activities are uncoupled, intermediates states proliferate and persist over a broader temporal and/or spatial window. Coordinated morphogen dynamics compress the transitional window, yielding an abrupt cell-state transition. The pink bar in wildtype indicates the spatial field of SHH activation. The dotted line represents the spatial profile of Wnt activity observed at E13.5 prior to SHH expression, modeled here as a baseline gradient. Data as mean± SEM; \*\**P*<0.01, \*\*\**P*<0.001, \*\*\*\**P*<0.0001, one-way ANOVA; ns, not significant; scale bars, 50 µm.

GT analysis identified three trajectories corresponding to proliferation, lower dermal differentiation and DC differentiation, mirroring E14.5 wildtype dermis (Figs. 4a and 7e and Extended Data Fig. 8b). Notably, a distinct Exit trajectory was not detected. Within the DC trajectory, *Lef1* and *Ptch1* increased coordinately, and DC genes and *Cdkn1a* were induced synchronously at the terminal stage, coincident with quiescence (Fig. 7f and Extended Data Fig. 8c). Thus, coupling Wnt and SHH activity is sufficient to restore the temporal alignment of differentiation and cell-cycle exit.

Based on our findings in Actβcat mutants, we next asked whether loss of GLI3 chromatin occupancy also occurs when Wnt and SHH signaling are coupled. GLI3 CUT&RUN analysis in SHH OE dermis revealed global depletion of GLI3 binding (Fig. 7g and Extended Data Fig. 8d), indicating that GLI3 displacement is not an artifact of genetic stabilization of β-catenin. Loss of GLI3 binding was observed at cell cycle regulatory genes (*Mxd4* and *Efna5*) associated with active histone marks (Extended Data Fig. 8e). In contrast to Actβcat mutants, SHH target genes (*Ptch1*, *Gli1*) were transcriptionally activated, accompanied by increased H3K4Me3 and H3K27Ac signal. Because SHH signaling converts cleaved GLI3 repressor into full-length GLI activator, both detectable by anti-Flag immunoprecipitation, the observed loss of GLI3 occupancy suggests replacement by other GLI activators at SHH target loci (Fig. 7h). These data indicate that GLI3 eviction both derepresses cell-cycle exit genes and may also prime SHH-dependent target loci for activation.

To test whether graded Wnt and SHH inputs are sufficient to coordinate these processes outside of the *in vivo* tissue context, we reconstituted the system *in vitro* by using E13.5 PDGFRαH2BGFP dermal cells competent to respond to both pathways^45^. These cells showed dose-dependent gene responses to the Wnt agonist (CHIR99021) or the Smoothened agonist (SAG) (Extended Data Fig. 9a). SAG enhanced *Lef1* expression in CHIR-treated cells in a manner dependent on baseline Wnt activity (Extended Data Fig. 9b). High doses of CHIR alone induced *Cdkn1a*, whereas combined CHIR and SAG treatment induced DC genes, including *Gal* and *Sox18* (Extended Data Fig. 9c,d).

Finally, to assess how signal coupling affects DC morphogenesis, E13.5 dermal cells were seeded onto a collagen-based 3D matrix with varying doses of CHIR and/or SAG (Fig. 7i and Extended Data Fig. 9e). High-dose CHIR (5 μM) combined with SAG (50 nM) produced large dermal clusters that expressed Robo2, a gene expressed in DC cells that regulates cell migration and tissue morphogenesis^46, 47^. These clusters showed low proliferation, whereas other conditions showed higher proportions of Ki67+ cells. Thus, coupling high Wnt and SHH activity is sufficient to align molecular differentiation with cell-cycle exit *in vitro*.

Collectively, these results indicate that the pre-DC–to-DC transition comprises separable biological processes whose timing can be differentially coordinated by Wnt and SHH. These relationships are summarized in 7j.

## Discussion

In this study, we identify a mechanism by which interacting morphogen signals generate sharp cell-type boundaries by coordinating the timing of separable components of a cell-state transition. During dermal condensate (DC) formation, we show that Sonic Hedgehog (SHH) acts on Wnt-primed cells to elevate Wnt activity, which is sufficient to induce cell-cycle exit through eviction of the transcriptional repressor GLI3 from chromatin, while SHH concurrently initiates the differentiation program. By aligning differentiation with cell-cycle exit, this coordination prevents proliferative expansion of intermediate states, causing transitional cells to remain sparse and difficult to detect. In this way, boundary sharpness reflects regulated temporal compression of a developmental transition, rather than the absence of intermediate cell states. While sharp transitions can also arise from nonlinear ligand diffusion, our findings reveal a distinct mechanism in which interactions between morphogen pathways synchronize concurrent transcriptional programs.

These findings revisit classic interpretations of the French flag problem, which emphasized fixed concentration thresholds of a polarized morphogen as primary determinants of spatial patterning^2–6, 48^. Our data instead indicate that during DC specification, morphogen thresholds function as triggers that initiate a committed developmental progression. In this context, SHH gates entry into the DC differentiation program, whereas Wnt activity shapes the timing and synchrony of its execution, establishing a division of labor between permissive and kinetic morphogen inputs. Abrupt boundaries therefore emerge not from static positional decoding, but from coordinated progression through a shared cell-state transition, providing a framework for understanding how gradual signaling changes can be converted into abrupt cell-state transitions.

Although tight coupling of Wnt and SHH activities generates a sharp and synchronous transition during DC formation, variation in the timing, strength, or spatial alignment of interacting morphogen gradients may support alternative modes of tissue organization. In systems containing transit-amplifying populations, differentiation programs can proceed without immediate cell-cycle exit, expanding intermediate states and producing graded or continuous boundaries^7, 49, 50^. Our findings suggest that the duration and apparent abundance of transitional states are tunable outcomes of morphogen coordination, providing a general framework for how interacting signals regulate the emergence of distinct tissue boundaries tissues (Fig. 7j).

How SHH signaling elevates Wnt activity is unlikely to involve direct transcriptional activation of Wnt target genes by GLI factors. Instead, our data indicate that elevated Wnt activity is associated with loss of GLI3 chromatin occupancy at loci involved in cell-cycle regulation.

Although GLI3 eviction from chromatin does not induce large-scale chromatin architectural reorganization, it coincides with SHH-dependent activation of differentiation genes, suggesting that chromatin-level cross-regulation between Wnt and SHH pathways synchronizes cell-cycle exit with molecular differentiation.

At the tissue level, this SHH-dependent enhancement of Wnt responsiveness shapes how transitions unfold in space and time. Prior to SHH signaling, dermal cells occupy positions along a Wnt-defined continuum, reflecting differences in transcriptional state. These differences encode competence – how rapidly and synchronously cells traverse the transition once SHH is engaged rather than pre-specified fates. Cells with higher Wnt activity reach cell-cycle exit more rapidly upon SHH activation, whereas cells with lower Wnt activity transition more slowly, expanding intermediate states. In this view, SHH synchronizes progression across a pre-patterned landscape, compressing developmental time locally and producing sharp boundaries without altering the sequence of states traversed.

Finally, by focusing on an early dermal transition, this work establishes a tractable mammalian system for investigating how interacting morphogen gradients regulate the timing of concurrent biological processes. Integrating *in vivo* genetics with trajectory-based analysis enables resolution of overlapping differentiation programs that are otherwise difficult to detect, revealing how dynamic signaling interactions shape discrete cell-state transitions^34^.

## Methods

### Tamoxifen induction of mice

Embryos were staged as days post coitum, with embryonic day E0.5 considered as noon of the day a vaginal plug was detected after overnight mating. Pregnant dams were given one dose of Tamoxifen dissolved in corn oil (20mg/ml, Sigma) at 40-60 µg/gm body weight by oral gavage at E11.5 as indicated.

### Mice details

*Axin2CreER* ^51^ mice were bred to *Rosa-LSLSmoM2YFP* ^52^, *β-catenin*^fl(Ex3*/+*^ ^53^ and *Gli3^fl/fl^* ^54^ mice. *K14Cre* ^55^ mice were bred to *Shh* Overexpressor (Wang et al. 2010) and *Wntless^fl/^*^fl56^mice. *Rosa-Flag-Gli3YFP* ^39^ mice were bred to *Axin2CreER* and *β-catenin*^fl(Ex3*/+*^ mice to enable detection of GLI3-Flag using CUT&RUN. *R26Fucci2aR* ^57^ reporter mice were used to visualize cell cycle phase. *PDGFRaH2BGFP*^45^ mice were used culture embryonic dermal cells. *Sox2eGFP*^58^ mice were used to visualize DC cells in embryonic skin whole mounts. A random population of both male and female mice were used for all experiments. All procedures involving animal subjects were performed under the approval of the Institutional Animal Care and Use Committee of the Yale School of Medicine.

### EdU Incorporation Assay

To assess active proliferation, EdU was administered to pregnant mice intraperitoneally (25 µg/gm) and embryos were harvested after 1.5 hour. For pulse-chase experiments, 25 µg/gm EdU was administered to pregnant mice at indicated times and embryos were harvested at later times. EdU incorporation was assessed using the Click-it EdU Imaging kit Alexa 555 or Alexa 488 (Life technologies, c10338) according to manufacturer’s instructions. Briefly, skin explants were treated with a mixture of 1X Click-iT reaction buffer, CuSO_4_, Alexa Fluor azide dye and 1X reaction buffer additive, all provided in the Click-it EdU Imaging Kit, for 30 minutes at RT before washing in PBS.

### Histology

10% formalin-fixed paraffin embedded (FFPE) whole embryos were used for histological analysis. FFPE embryos were sectioned at 10 µm thickness for FISH.

### Whole mount immunofluorescence

Dorsolateral skin from embryos were microdissected and placed on nucleopore filters (VWR, WHA 800281) and fixed overnight with 4%PFA at 4° C. Skin explants were blocked for 6 hours in 5% normal donkey serum, 1% bovine serum albumin and 0.5% Triton-X100 at RT. Explants were incubated overnight at 4° C in the following primary antibodies: rabbit anti-Sox2 (1:400, Abcam, ab97959), chicken anti-GFP (1:500, Abcam, ab13970), rabbit anti-RFP (1:500, Rockland, 600-401-379). Explants were then washed in 0.2% Tween20/PBS for 4-6 hours on a rotator and then incubated with the following secondary antibodies: Alexa Fluor 568-donkey anti-rabbit (1:200, Invitrogen, A10042), Alexa Fluor 488-donkey anti-rabbit (1:200, Invitrogen, A21206), Alexa Fluor 488-goat anti-chicken (1:200, Invitrogen, A11039) overnight at RT. Washes were carried out for 6 hours with 0.2% Tween20/PBS. Hoechst (1:750 in PBS) was used for 45 min before mounting on slides with SlowFade Gold Antifade Mountant (Invitrogen, S36936).

### *In-situ* hybridization

The RNAscope Multiplex Fluorescent Detection Kit v2 (ACDBio, 323110) was used for single-molecule fluorescence in situ hybridization (FISH) according to the manufacturer’s protocol. Briefly, sections were deparaffinized and permeabilized with hydrogen peroxide followed by antigen retrieval and protease treatment before probe hybridization. After hybridization, amplification and probe detection were done using the Amp 1–3 reagents. Probe channels were targeted using the provided HRP-C1-3 reagents and tyramide signal amplification VIVID fluorophores—650 and 570 (ACD biotechne, 323271). EdU staining was done using the Click-it EdU Imaging Kit Alexa 488 (Life Technologies, c10338) according to the manufacturer’s instructions. Nuclear counter-stain was done using Hoechst 33342 (Invitrogen, H3570) before mounting with SlowFade Mountant. RNA scope probes used (ACDBio)—Mm-Lef1 (441861), Mm-Ptch1 (402811), Mm-Dkk1 (402521), Mm-Sox2 (401041), Mm-Gal (400961), Mm-Gli3 (400961) and Mm-Trp53inp1 (1161531).

### Microscopy

FISH paraffin-embedded images were acquired using the Leica Stellaris 8 DMi8 confocal microscope with a ×40 oil immersion (Numerical Aperture 1.3) objective lens, scanned at 5 µm thickness, 1, 024 × 1, 024 pixel width, 400Hz. Whole mount explants were imaged in 3 dimensions using the LaVision TriM Scope II (LaVision Biotec) microscope equipped with a Chameleon Vision II (Coherent) two-photon laser (810 -1000 nm) to acquire *z*-stack images ranging from 50-120 μm (2 μm serial optical sections) using a 20X water immersion lens (NA 1.0; Olympus), scanned with a field of view of 0.3–0.5 mm^2^ at 800 Hz.

### Single-cell dissociation

Embryonic dorsolateral/flank skin was micro-dissected and pooled after genotyping (3 embryos per condition) and dissociated into a single-cell suspension using 0.25% trypsin (Gibco, Life Technologies) for 14 minutes at 37° C. Single-cell suspensions were then stained with DAPI (Fisher Scientific, NBP2-31156) just prior to fluorescence-activated cell sorting.

### Fluorescence-activated cell sorting

DAPI-excluded live skin cells were sorted on a BD FACS Aria II (Biosciences) sorter with a 100 µm nozzle. Cells were sorted in bulk and submitted for 10X Genomics library preparation at 0.75-1.0x10^6^/mL concentration in 4% FCS/PBS solution.

### Single-cell RNA sequencing and library preparation

Chromium Single cell 3’ Library and Gel Bead Kit v2 Chromium Single Cell 3′ Library & Gel Bead Kit v2 (PN-120237), Chromium Single Cell 3′ Chip kit v2 (PN-120236) and Chromium i7 Multiplex Kit (PN-120262) were used according to the manufacturer’s instructions in the Chromium Single Cell 3′ Reagents Kits V2 User Guide. After cDNA libraries were created, they were subjected to Novaseq 6000 (Illumina) sequencing.

### 10X scRNA-seq data pipeline to matrix

The transcriptomes of live single cells from E13.5 or E14.5 mouse skin samples under different biological conditions were sequenced. Raw 10X sequencing data was processed into a matrix employing the standard 10X CellRanger pipeline. Briefly, base call files were fastq format which were aligned to the mm10 reference genome followed by nUMI and barcode counting, constructing the nUMI count matrices. nUMI matrices were filtered, centered and normalized using Seurat@1. Briefly, for each original run condition (e.g., E13.5 wildtype), cells with > 1,000 detected genes, mitochondrial content < 25%, and ribosomal content > 5% (calculated via PercentageFeatureSet in Seurat) were retained. Doublets were identified and removed using scDblFinder@2. Data were then log-normalized by total UMI count.

### Principal Component Calculation and UMAP Visualization

Principal components were computed using Seurat’s RunPCA on the top 2,000 variable genes. The number of PCs retained was determined from the elbow plot@7, and used for downstream analysis, including dimensionality reduction for visualization and clustering.

### Cell type specification

For each original run, UMAP was performed on log-normalized, centered, and scaled UMI count matrices using the retained PCs. Unsupervised clustering was then conducted using Seurat’s FindClusters function with a resolution of 0.5. Clusters expressing Col1a1 were defined as dermal, while clusters expressing Lef1 or Dkk1 were defined as upper-dermal.

### Gene trajectory inference

We applied GeneTrajectory@3 to each sample to identify multiple, independent gene dynamic processes. For each sample, we followed the recommended guidelines. We selected the top 2,000 highly variable genes (identified using Seurat’s FindVariableFeatures) that were expressed in >1% and <50% of cells as input. The top 20PCs were used to compute a diffusion map, and the top 10 diffusion components were used to construct a cell k-nearest neighbor graph (k = 10).

The number of gene trajectories and the time steps for gene inclusion within each trajectory were determined interactively by inspecting the top three or more gene embeddings. For the samples analyzed in this study, we identified the following gene trajectories and associated time steps (t):

- E14.5 wild type: 3 trajectories (t = 13, 10, 5)
- E13.5 SmoM2: 4 trajectories (t = 1, 3, 2, 1); control: 3 trajectories (t = 3, 2, 2)
- E13.5 TG: 3 trajectories (t = 6, 3, 3); control: 3 trajectories (t = 5, 3, 3)
- E13.5 EX3: 4 trajectories (t = 3, 3, 2, 2); control: 3 trajectories (t = 1, 3, 3)
- E13.5 Gli3-KO: 4 trajectories (t = 8, 7, 6, 1); control: 3 trajectories (t = 4, 3, 1)

### Gene bins and cell stages

Following GeneTrajectory guidelines, each trajectory from mutant samples was divided into 5 or 6 gene bins. For each bin, a gene bin score was computed for both mutant and control samples to compare gene expression dynamics across the cell embedding. Cell involved in each trajectory were stratified into the same number of stages, based on whether they expressed more than 50% of the genes in the corresponding gene bin. Cells assigned to multiple stages were resolved by assigning them to the stage closer to the terminal end of the trajectory (i.e., farther from the branching point).

### Exit GT Score

Exit GT scores for each sample were calculated using Seurat’s AddModuleScore, based on all genes from the exit gene trajectory identified in E13.5 SmoM2.

### Pathway enrichment analysis

Reactome pathway enrichment analysis was performed using ReactomePA@4, with pathways showing p < 0.05 considered significantly enriched.

### Cell cycle estimation of scRNA-seq data

We employed Seurat’s cell cycle scoring ^59^. Briefly, averaged relative expression of those cell cycle related genes were used to calculate G2/M and S scores, which were used for binning cells into G2/M, S and G1/G0 bins.

### Data merge and normalization

For comparative analysis, we merged upper-dermal cell matrices from each pair of conditions separately (i.e. E13.5 control and Gli3-KO), followed by standard preprocessing and normalization using the workflow provided by Seurat.

### Differential abundance and Differential gene expression analysis

Differential abundant cells were identified using DA-seq@6, with a DA score threshold > 0.8. Differential gene expression analysis between DA cells from E13.5 Gli3-KO and all other upper-dermal cells from both E13.5 Gli3-KO and control was performed using Seurat’s FindMarkers without prefiltering. Significant DEGs for visualization were defined using a combined threshold of padj < 1e-5 and log2FC > 0.23.

### Integrated snATAC+RNA-seq Multi-omic Analysis

Single-cell RNA-seq (scRNA-seq) and single-cell ATAC-seq (scATAC-seq) data from E14.5 K14Cre;SHH^fl/fl^ and paired control embryonic skin were processed and integrated using the Seurat and Signac packages in R. The scRNA-seq data was normalized using SCTransform. For the scATAC-seq component, dimensionality reduction was performed using Latent Semantic Indexing (LSI). To integrate the modalities, we applied Weighted Nearest Neighbor (WNN) analysis, constructing a joint neighbor graph that represents the weighted combination of RNA and ATAC modalities. A joint UMAP was generated from the WNN graph for visualization. Transcription factor motif accessibilities were calculated using chromVAR.

### CUT&RUN experiments

CUT&RUN was performed with the Epicypher CUTANA™ ChIC/CUT&RUN Kit (14-1048) according to the manufacturer’s specifications with the following conditions: 500,000 cells per sample were prepared first and nuclei were extracted according to the Epicypher CUT&RUN Manual Appendix. Nuclei were incubated with activated ConA beads. The following antibodies (0.5 ug of antibody per reaction) were used:

IgG Control antibody: CUTANA™ Kit Rabbit IgG CUT&RUN Negative Control Antibody, EpiCypher Rabbit anti-H3K27Ac antibody, Millipore Mouse anti-Flag antibody, Antibodies-online Rabbit anti-CTNNB1 antibody.

Libraries were prepared using the Epicypher CUTANA™ CUT&RUN Library Prep Kit (14-1002) according to the manufacturer’s specifications. Library quality was analyzed using Agilent Tapestation D1000 High Sensitivity Tapes (#5067-5584). Sequencing was performed with 5 million 150bp paired-end reads per sample on an Illumina NovaSeq 6000.

### CUT&RUN data analysis

CUT&RUN reads were analyzed for quality using FastQC (https://www.bioinformatics.babraham.ac.uk/projects/fastqc/) and adaptors trimmed using Trimmomatic. Reads were then aligned to mm9 by Bowtie2 version 2.3.1. SAMTools version 1.11 was then used to convert to BAM format, index, isolate uniquely mapped paired reads, and remove duplicates. MACS2 version 2.2.8 was used to call peaks with IgG as input control. Read counts across genomic intervals and peak visualization were performed using deepTools version 3.5 and bigwig files were visualized using the Integrative Genomics Viewer. Deseq2 was used to identify differentially bound peaks between conditions, DiffBind used to plot heatmaps of accessible/bound regions and HOMER to identify *de novo* motifs most enriched in the differentially bound regions and annotate associated genes.

### Micro-C Experiments

Micro-C was performed using the Dovetail® Micro-C Kit (Cantata Bio, PN 21006E) according to the manufacturer’s protocol using 1 million cells per reaction. E13.5 mouse dorsolateral dermis was microdissected, dissociated, and crosslinked with formaldehyde, followed by quenching and nuclei isolation. Chromatin was digested with micrococcal nuclease, end-repaired, and proximity ligated to capture nucleosome-level chromatin contacts. Reverse crosslinking and DNA purification were performed following kit specifications. Libraries were prepared using the Dovetail® Library Module for Illumina (PN 25004E) and indexed with the manufacturer-provided i5/i7 primer sets. Library quality and fragment size distribution were assessed using the Agilent TapeStation High Sensitivity D5000 assay (Agilent #5067-5592**, #**5067-5593**, #**5067-5594). Final libraries were sequenced on an Illumina NovaSeq 6000 platform to a depth of 300 million 150-bp paired-end reads per biological replicate.

### Micro-C data analysis

Micro-C reads were aligned to the *mm10* genome using BWA-MEM with proximity-ligation–optimized settings, and ligation junctions were identified, sorted, and deduplicated using the Pairtools pipeline to generate uniquely mapped, non-duplicate valid pairs. Valid pairs were converted into 5-kb contact matrices using cooler (v0.10.4), balanced by iterative correction, and aggregated into multi-resolution .mcool files for downstream analysis. Chromatin loops were identified using Mustache (v1.2.0), a multi-scale loop-calling algorithm that applies scale-space representation to detect interaction peaks across resolutions in high-resolution Micro-C maps. Loop anchors were intersected with CUT&RUN peaks using Bedtools (v2.27.1) to annotate interactions. Genome-wide A/B chromatin compartments were computed at 40-kb resolution using FAN-C (v0.9.28), which generated Pearson correlation matrices and extracted the first principal component to assign A/B compartment identity. Processed matrices, loops, and eigenvector tracks were visualized using FAN-C.

### Western blot

Cells were lysed in RIPA buffer (Thermo Fisher Scientific, Cat# 89900) supplemented with protease inhibitors (Thermo Fisher Scientific, Cat# 87786) for 30 minutes at 4 °C. The lysates were clarified by centrifugation at 10,000 × g for 20 minutes at 4 °C, and protein concentrations were determined using Bradford Protein Assay Kit (Bio-Rad, Cat# 5000006). Equal amounts of protein (10 µg) were denatured at 95 °C for 5 minutes and loaded onto 4–12% precast polyacrylamide gels (Bio-Rad, Cat# 3450125) for electrophoresis. Proteins were transferred to PVDF membranes (Thermo Fisher Scientific, Cat# IB24001) using an iBlot2 transfer device (Invitrogen, Cat# IB21001). Membranes were blocked with 5% BSA (vol/vol) in 1× TBST (AmericanBio, Cat# AB14330-01000) at room temperature, incubated with primary antibodies overnight at 4 °C, washed three times with TBST for 5 minutes each, incubated with HRP-conjugated secondary antibodies for 1 hour at room temperature, and washed again three times with TBST.

Membranes were developed using SuperSignal West Pico PLUS Chemiluminescent Substrate (Thermo Fisher Scientific, Cat# 34577) and imaged on a ChemiDoc imaging system (Bio-Rad). The following antibodies were used: mouse anti-Flag (Millipore, Cat# F1804), rabbit anti-Actin (Cell Signaling Technology, Cat# 4970), HRP-conjugated goat anti-rabbit IgG (Cell Signaling Technology, Cat# 7074), and HRP-conjugated goat anti-mouse IgG (Bio-Rad, Cat# STAR207P, 1:10,000).

### Dermal cell culture

E13.5 PDGFRαH2BGFP skin cells were sorted based on their GFP status. GFP+ cells were collected and pelleted (300 × g, 10 min, 4°C), resuspended at 2 × 10⁶ cells/mL in complete growth medium (DMEM + 10% FBS, Penicillin-Streptomycin, and L-Glutamine), and viability was assessed by trypan blue exclusion. Cells were plated at 3 × 10⁶ cells/well in 6-well plates in 2 mL total volume. For chemical perturbation assays, cells were mixed 1:1 with a 2× agonist solution. For example, 5 µM CHIR99021 (Sigma, Cat# SML1046) was prepared in DMSO and diluted to a working concentration of 5 µM in culture. Similarly, SAG (Sigma, Cat# SML1314) was prepared as a 50 µM stock and diluted as needed. Media with agonist was used fresh or stored at 4°C for up to two days. Cells were cultured in 48-72 hours. For 3D culture, 3 × 10^5^ cells were seeded onto each well of a μ-slide 8 Well Glass Bottom chamber (ibidi, Cat#80807) which has been precoated by 1.5mg/mL Collagen Type I (Corning, Cat#354236). Collagen Type I was neutralized by manufacture instructions and 200uL was added to each well 1 hour prior to seeding the cells for solidification.

### Quantitative PCR

Total RNA was extracted from cultured cells using the RNeasy Mini Kit (Qiagen, Cat# 74134) according to the manufacturer’s protocol. Complementary DNA (cDNA) was synthesized from 500 ng–1 µg of total RNA using the SuperScript™ III First-Strand Synthesis System (Invitrogen, Cat# 18080051) with random hexamer primers. Quantitative PCR was performed using the PowerUp™ SYBR™ Green Master Mix (Thermo Fisher Scientific, Cat# 4364344) on a Viia7 Real-Time PCR System (Applied Biosystems) in 384-well optical plates. Reactions were run in technical triplicates using gene-specific primers, and threshold cycle (Ct) values were normalized to Gapdh as a housekeeping gene. Relative expression was calculated using the ΔΔCt method.

### 3D culture immunofluorescence

Cultured cells with collagen were fixed for 25 minutes with 4%PFA at room temperature. They were blocked for 2-4 hours in 5% normal donkey serum, 1% bovine serum albumin and 0.2% Triton-X100 at RT. Cells were incubated overnight at 4° C in the following primary antibodies: rat anti-Ki67 (1:200, Invitrogen, 14-5698-82), rabbit anti-Robo2 (1:200, Cell signaling, 45568). Cells were then washed in 1x PBS for 4-6 hours and then incubated with the following secondary antibodies: Alexa Fluor 568-goat anti-rat (1:200, Invitrogen, A11077) and Alexa Fluor 647-donkey anti-rabbit (1:200, Invitrogen, A31573) overnight at 4° C. Washes were carried out for 6 hours with 1x PBS. Hoechst (1:750 in PBS) was used for 45 min before washing in 1x PBS.

### Image analysis

Raw image stacks acquired from whole-mount explants were imported into Fiji or Adobe Photoshop for analysis. ggplot2 and cowplot R libraries were used for graphical representation of the scRNA-seq data. Diffusion maps were generated as described above.

### Quantification

For quantification of DC density, 3-dimensional whole-mount tiled mosaics of skin explants with a 1000 x 1000 µm field of view and a z-depth of 60-100 µm were used for n=3-4 embryos. For volumetric quantification of DC cell number, EdU+ or Tom+ percentage of DC or UD, 3-dimensional whole mount mosaics stained with Sox2 and EdU or RFP were used to manually count positive cells, using ImageJ (Fiji) software using planar (XY) and orthogonal (YZ, XZ) views. For volumetric quantification of cell cycle phases, 3-dimensional whole mount mosaics stained with markers for G0/G1 cells (based on Fucci2 reporter) were done manually by counting nuclear positive cells using ImageJ (Fiji) software. For quantification of upper dermal cells, a distance of 10 um below the epidermis was considered upper dermis. For quantification of the peri-DC region by whole-mount, cells two cell layers out from the Sox2+ DC were counted. For quantification based on FISH, cells with 4-5 dots were considered positive (according to the RNAScope manufacturer’s instructions) and sections from a total of n=4 different embryos were examined. To measure RNA expression levels, H-scores were calculated according to ACDBio manufacturer’s instructions: a cell with 0 dot is scored 0, 1-3 dots score 1, 4-9 dots score 2, 10-15 dots and/or less than 10% clustered dots score 3, and more than 15 dots and/or more than 10% clustered dots score 4; then the final H-score of a given cell type A is calculated by summing the (% cells scored B within all cells in A)*B for score B in 0-4. For quantification of upper dermal cells, a distance of 10-20 μm below the epidermis was considered as upper dermis. For quantification of cells in the peri-DC region *Dkk1*+*Sox2*-was considered.

### Statistical analysis

All statistical values are expressed as mean ± SEM. An unpaired Student’s *t*-test was used to analyze data sets with two groups and \**P* <0.05 to, \*\**P*<0.01, \*\*\**P*< 0.001 and \*\*\*\**P*<0.00001 indicated a significant difference. When comparing more than two groups, *P* values were determined by one-way ANOVA with Tukey’s HSD test performed as the post hoc analysis. Statistical calculations were performed using Prism software package (GraphPad). For categorical data, a chi-square test was employed to estimate a *P* value.

No statistical methods were used to predetermine <u>sample size</u>. The experiments were not randomized. The investigators were not blinded to allocation during experiments and outcome assessment.

## Data and Code Availability

Genomic datasets used in this study will be deposited in NCBI Gene Expression Omnibus database and made publicly available as of the date of publication or with a token upon request.

## Acknowledgements

We thank D. Gay, M. Ito, J. Levinsohn, and R. Atit for review and discussion of this manuscript. We thank J. Wang and K. Sumigray for generously sharing their Stellaris equipment and help with imaging troubleshooting. Inducible SHH ligand transgenic mice were kindly provided by David Wang. This work was supported by National Institutes of Health grants R01AR076420 (P.M.) and LEO Foundation LF-OC-23-001347 (P.M.).

## Author Contributions

Conceptualization, P.M., Y.K., R.L., Y.J., S.P.; Methodology, P.M., R.L., Y.K., Y.J., S.P., E.B., S.V., H.L.; Software, R.L., Y.K.; Resources, M.T., K.P.; Investigation, P.M., R.L., Y.J., S.P. H.L, E.B., S.V.; Writing – Original Draft, P.M.; Writing – Review & Editing, P.M., R.L., Y.J., S.P., T.X., R.D., Y.K., C.L., S.L., E.B.; Supervision, P.M., Y.K.; Funding Acquisition, P.M.

## References

1. Wolpert, L. Positional information and the spatial pattern of cellular differentiation. J Theor Biol 25, 1–47 (1969).

2. Benzinger, D. & Briscoe, J. Investigating morphogen and patterning dynamics with optogenetic control of morphogen production. Dev Cell 60, 3421–3430 e3426 (2025).

3. Driever, W. & Nusslein-Volhard, C. The bicoid protein determines position in the Drosophila embryo in a concentration-dependent manner. Cell 54, 95–104 (1988).

4. Greenfeld, H., Lin, J. & Mullins, M.C. The BMP signaling gradient is interpreted through concentration thresholds in dorsal-ventral axial patterning. PLoS Biol 19, e3001059 (2021).

5. Roelink, H. et al. Floor plate and motor neuron induction by different concentrations of the amino-terminal cleavage product of sonic hedgehog autoproteolysis. Cell 81, 445–455 (1995).

6. Teague, S. et al. Time-integrated BMP signaling determines fate in a stem cell model for early human development. Nat Commun 15, 1471 (2024).

7. Morita, R. et al. Tracing the origin of hair follicle stem cells. Nature 594, 547–552 (2021).

8. Ouspenskaia, T., Matos, I., Mertz, A.F., Fiore, V.F. & Fuchs, E. WNT-SHH Antagonism Specifies and Expands Stem Cells prior to Niche Formation. Cell 164, 156–169 (2016).

9. Rompolas, P. et al. Live imaging of stem cell and progeny behaviour in physiological hair-follicle regeneration. Nature 487, 496–499 (2012).

10. Zhang, C. et al. Escape of hair follicle stem cells causes stem cell exhaustion during aging. Nat Aging 1, 889–903 (2021).

11. Leybova, L. et al. Radially patterned morphogenesis of murine hair follicle placodes ensures robust epithelial budding. Dev Cell 59, 3272–3289 e3275 (2024).

12. Driskell, R.R. et al. Distinct fibroblast lineages determine dermal architecture in skin development and repair. Nature 504, 277–281 (2013).

13. Myung, P., Andl, T. & Atit, R. The origins of skin diversity: lessons from dermal fibroblasts. Development 149 (2022).

14. Chuong, C.M., Widelitz, R.B., Ting-Berreth, S. & Jiang, T.X. Early events during avian skin appendage regeneration: dependence on epithelial-mesenchymal interaction and order of molecular reappearance. J Invest Dermatol 107, 639–646 (1996).

15. Chase, H.B., Rauch, R. & Smith, V.W. Critical stages of hair development and pigmentation in the mouse. Physiol Zool 24, 1–8 (1951).

16. Gupta, K. et al. Single-Cell Analysis Reveals a Hair Follicle Dermal Niche Molecular Differentiation Trajectory that Begins Prior to Morphogenesis. Dev Cell 48, 17–31 e16 (2019).

17. Hardy, M.H. The secret life of the hair follicle. Trends Genet 8, 55–61 (1992).

18. Millar, S.E. et al. WNT signaling in the control of hair growth and structure. Dev Biol 207, 133–149 (1999).

19. Paus, R. et al. A comprehensive guide for the recognition and classification of distinct stages of hair follicle morphogenesis. J Invest Dermatol 113, 523–532 (1999).

20. Saxena, N., Mok, K.W. & Rendl, M. An updated classification of hair follicle morphogenesis. Exp Dermatol 28, 332–344 (2019).

21. Woo, W.M., Zhen, H.H. & Oro, A.E. Shh maintains dermal papilla identity and hair morphogenesis via a Noggin-Shh regulatory loop. Genes Dev 26, 1235–1246 (2012).

22. Chen, D., Jarrell, A., Guo, C., Lang, R. & Atit, R. Dermal beta-catenin activity in response to epidermal Wnt ligands is required for fibroblast proliferation and hair follicle initiation. Development 139, 1522–1533 (2012).

23. Noramly, S., Freeman, A. & Morgan, B.A. beta-catenin signaling can initiate feather bud development. Development 126, 3509–3521 (1999).

24. Chiang, C. et al. Essential role for Sonic hedgehog during hair follicle morphogenesis. Dev Biol 205, 1–9 (1999).

25. St-Jacques, B. et al. Sonic hedgehog signaling is essential for hair development. Curr Biol 8, 1058–1068 (1998).

26. Gritli-Linde, A. et al. Abnormal hair development and apparent follicular transformation to mammary gland in the absence of hedgehog signaling. Dev Cell 12, 99–112 (2007).

27. Qu, R. et al. Decomposing a deterministic path to mesenchymal niche formation by two intersecting morphogen gradients. Dev Cell 57, 1053–1067 e1055 (2022).

28. Xu, Z. et al. Embryonic attenuated Wnt/beta-catenin signaling defines niche location and long-term stem cell fate in hair follicle. Elife 4, e10567 (2015).

29. Ben-Zvi, D., Shilo, B.Z., Fainsod, A. & Barkai, N. Scaling of the BMP activation gradient in Xenopus embryos. Nature 453, 1205–1211 (2008).

30. Dekanty, A. & Milan, M. The interplay between morphogens and tissue growth. EMBO Rep 12, 1003–1010 (2011).

31. Gregor, T., Wieschaus, E.F., McGregor, A.P., Bialek, W. & Tank, D.W. Stability and nuclear dynamics of the bicoid morphogen gradient. Cell 130, 141–152 (2007).

32. Kicheva, A., Bollenbach, T., Wartlick, O., Julicher, F. & Gonzalez-Gaitan, M. Investigating the principles of morphogen gradient formation: from tissues to cells. Curr Opin Genet Dev 22, 527–532 (2012).

33. Lander, A.D., Nie, Q. & Wan, F.Y. Do morphogen gradients arise by diffusion? Dev Cell 2, 785–796 (2002).

34. Qu, R. et al. Gene trajectory inference for single-cell data by optimal transport metrics. Nat Biotechnol (2024).

35. Biggs, L.C. et al. Hair follicle dermal condensation forms via Fgf20 primed cell cycle exit, cell motility, and aggregation. Elife 7 (2018).

36. Zhang, Y. et al. Reciprocal requirements for EDA/EDAR/NF-kappaB and Wnt/beta-catenin signaling pathways in hair follicle induction. Dev Cell 17, 49–61 (2009).

37. Fu, J. & Hsu, W. Epidermal Wnt controls hair follicle induction by orchestrating dynamic signaling crosstalk between the epidermis and dermis. J Invest Dermatol 133, 890–898 (2013).

38. Alvarez-Medina, R., Cayuso, J., Okubo, T., Takada, S. & Marti, E. Wnt canonical pathway restricts graded Shh/Gli patterning activity through the regulation of Gli3 expression. Development 135, 237–247 (2008).

39. Vokes, S.A., Ji, H., Wong, W.H. & McMahon, A.P. A genome-scale analysis of the cis-regulatory circuitry underlying sonic hedgehog-mediated patterning of the mammalian limb. Genes Dev 22, 2651–2663 (2008).

40. te Welscher, P., et al. Progression of vertebrate limb development through SHH-mediated counteraction of GLI3. Science 298, 827–830 (2002).

41. Heinz, S. et al. Simple combinations of lineage-determining transcription factors prime cis-regulatory elements required for macrophage and B cell identities. Mol Cell 38, 576–589 (2010).

42. Lex, R.K. et al. GLI transcriptional repression is inert prior to Hedgehog pathway activation. Nat Commun 13, 808 (2022).

43. Phan, Q.M. et al. Lef1 expression in fibroblasts maintains developmental potential in adult skin to regenerate wounds. Elife 9 (2020).

44. Reddy, S. et al. Characterization of Wnt gene expression in developing and postnatal hair follicles and identification of Wnt5a as a target of Sonic hedgehog in hair follicle morphogenesis. Mech Dev 107, 69–82 (2001).

45. Hamilton, T.G., Klinghoffer, R.A., Corrin, P.D. & Soriano, P. Evolutionary divergence of platelet-derived growth factor alpha receptor signaling mechanisms. Mol Cell Biol 23, 4013–4025 (2003).

46. Guzman-Palma, P. et al. Slit/Robo Signaling Regulates Multiple Stages of the Development of the Drosophila Motion Detection System. Front Cell Dev Biol 9, 612645 (2021).

47. Goncalves, A.N., Correia-Pinto, J. & Nogueira-Silva, C. ROBO2 signaling in lung development regulates SOX2/SOX9 balance, branching morphogenesis and is dysregulated in nitrofen-induced congenital diaphragmatic hernia. Respir Res 21, 302 (2020).

48. Reyes, R., Lander, A.D. & Nahmad, M. Dynamic readout of the Hh gradient in the Drosophila wing disc reveals pattern-specific tradeoffs between robustness and precision. Elife 13 (2024).

49. Sun, Q. et al. Dedifferentiation maintains melanocyte stem cells in a dynamic niche. Nature 616, 774–782 (2023).

50. Myung, P. & Ito, M. Dissecting the bulge in hair regeneration. J Clin Invest 122, 448–454 (2012).

51. van Amerongen, R., Bowman, A.N. & Nusse, R. Developmental stage and time dictate the fate of Wnt/beta-catenin-responsive stem cells in the mammary gland. Cell Stem Cell 11, 387–400 (2012).

52. Jeong, J., Mao, J., Tenzen, T., Kottmann, A.H. & McMahon, A.P. Hedgehog signaling in the neural crest cells regulates the patterning and growth of facial primordia. Genes Dev 18, 937–951 (2004).

53. Harada, N. et al. Intestinal polyposis in mice with a dominant stable mutation of the beta-catenin gene. EMBO J 18, 5931–5942 (1999).

54. Blaess, S., Stephen, D. & Joyner, A.L. Gli3 coordinates three-dimensional patterning and growth of the tectum and cerebellum by integrating Shh and Fgf8 signaling. Development 135, 2093–2103 (2008).

55. Dassule, H.R., Lewis, P., Bei, M., Maas, R. & McMahon, A.P. Sonic hedgehog regulates growth and morphogenesis of the tooth. Development 127, 4775–4785 (2000).

56. Carpenter, A.C., Rao, S., Wells, J.M., Campbell, K. & Lang, R.A. Generation of mice with a conditional null allele for Wntless. Genesis 48, 554–558 (2010).

57. Abe, T. et al. Visualization of cell cycle in mouse embryos with Fucci2 reporter directed by Rosa26 promoter. Development 140, 237–246 (2013).

58. Arnold, K. et al. Sox2(+) adult stem and progenitor cells are important for tissue regeneration and survival of mice. Cell Stem Cell 9, 317–329 (2011).

59. Nestorowa, S. et al. A single-cell resolution map of mouse hematopoietic stem and progenitor cell differentiation. Blood 128, e20–31 (2016).

