## Extended Data Figures for "Sharp cell-type boundaries emerge from temporal coordination between morphogen signals"

### Extended Data Fig. 1

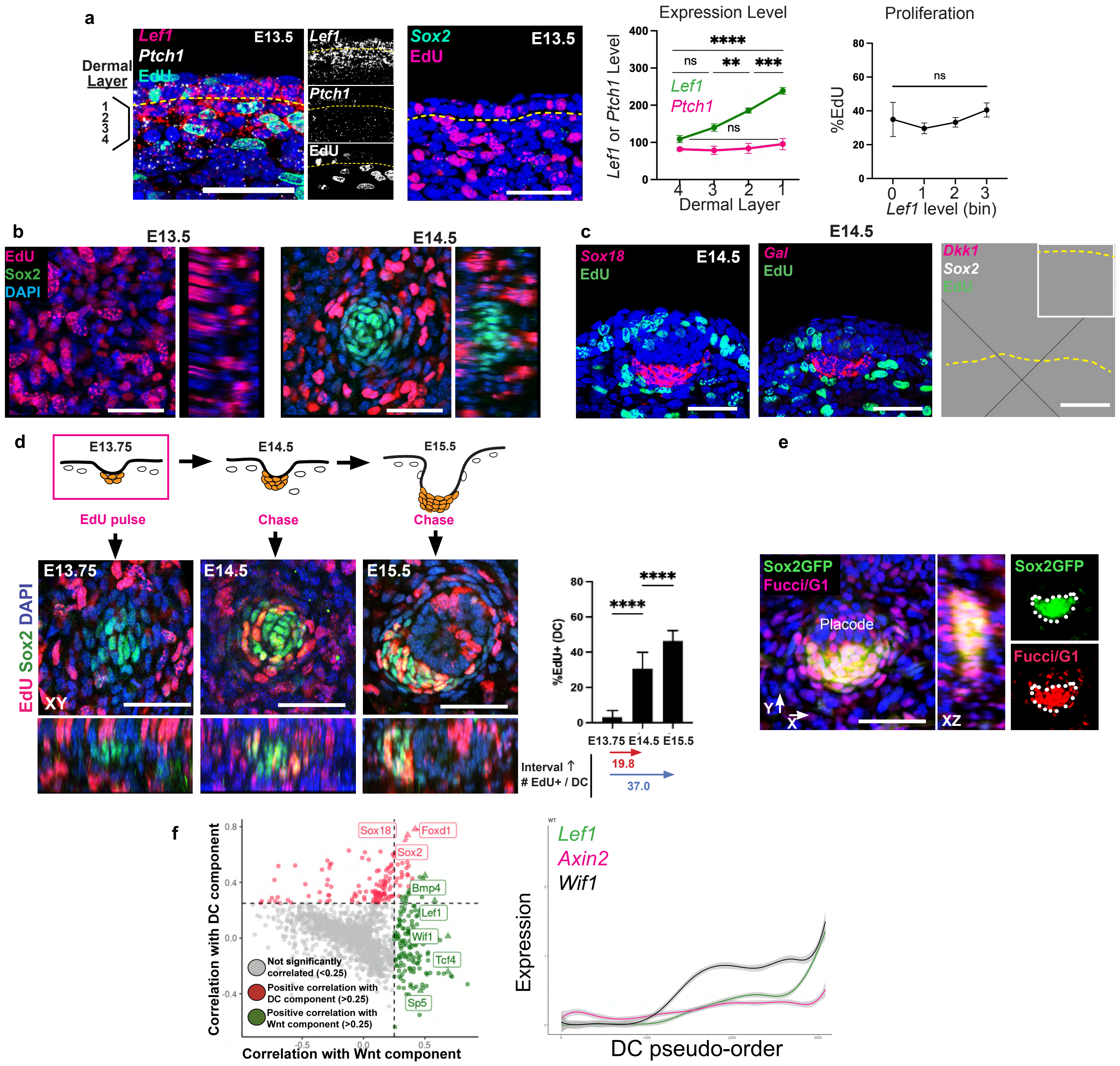


**Extended Data Fig. 1. Cell-cycle arrest and DC gene expression are temporally aligned at E14.5. a**, FISH image of E13.5 skin showing that *Lef1* levels decrease vertically across dermal layers before SHH; %EdU shows no correlation with *Lef1* levels. **b**, Whole mount images of E13.5 and E14.5 skin stained for Sox2 and EdU. **c,** FISH images showing cells that express DC markers, *Sox18, Gal, Sox2* lack EdU, while *Dkk1*+ peri-DC cells are proliferative (EdU+). **d,** Cartoon illustrating EdU pulse/chase labeling strategy from E13.75 to E15.5 to track proliferating DC progenitors. Corresponding whole mount images (below) showing EdU and Sox2 to track the transition from proliferation to quiescence. Corresponding quantification of EdU incorporation in DC cells. **e**, Whole mount image of skin stained for the cell cycle reporter Fucci2 (RFP) and Sox2eGFP, showing close overlap of stains. **f**, Correlation plot of E14.5 wildtype dermal scRNA-seq data, showing DC and Wnt target genes that correlate with *Sox2* and *Lef1*, respectively. Right, expression of *Lef1*, *Axin2* and *Wif1* as a function of DC cell pseudotime based on E14.5 scRNA-seq data. Data as mean± SEM; *P<0.05, **P<0.01, ***P<0.001, ****P<0.0001, one-way ANOVA; scale bars, 50 μm.

**Extended Data Fig. 2**

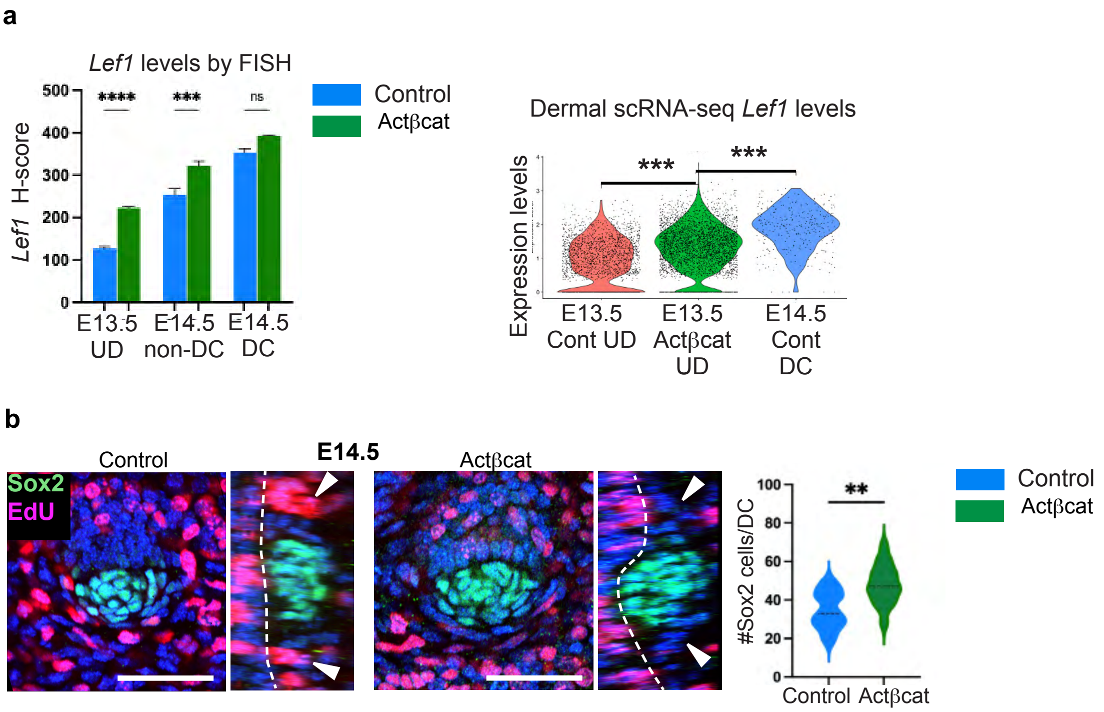


**Extended Data Fig. 2. β-catenin activation elevates dermal Wnt signaling and results in larger quiescent DCs at E14.5. a**, Left, FISH quantification of *Lef1* mRNA by cell type at E13.5 and E14.5 in control and Actβcat embryos. Right, *Lef1* expression based on scRNA-seq data of control or Actβcat mutant dermal cells (UD, upper dermis). **b**, Orthogonal whole mount images showing EdU and Sox2 stained skin from E14.5 control and Actβcat embryos. Right, quantification of Sox2+ cell number per DC by condition. Data as mean± SEM; *P<0.05, **P<0.01, ***P<0.001, ****P<0.0001, one-way ANOVA; scale bars, 50 μm.

### Extended Data Fig. 3

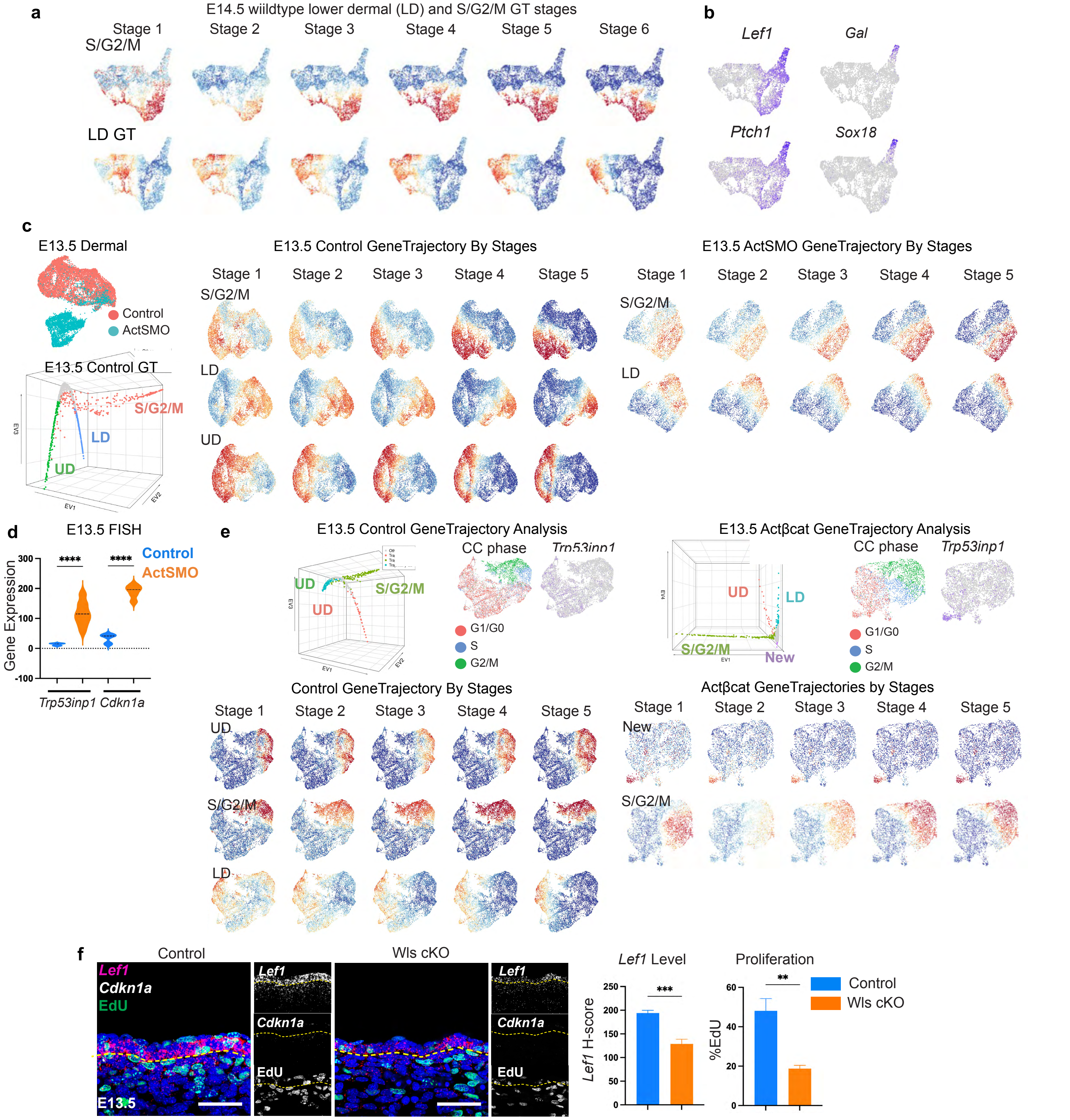


**Extended Data Fig. 3. GeneTrajectory method detects a gene program β-catenin activation elevates dermal Wnt signaling results in larger quiescent DCs at E14.5. a**, Orthogonal whole mount images showing EdU and Sox2 stained skin from E14.5 control and Actβcat embryos. Right, quantification of Sox2+ cell number per DC by condition. **b**, Left, FISH quantification of *Lef1* mRNA by cell type at E13.5 and E14.5 in control and Actβcat embryos. Right, *Lef1* expression based on scRNA-seq data of control or Actβcat mutant dermal cells (UD, upper dermis). Data as mean± SEM; *P<0.05, **P<0.01, ***P<0.001, ****P<0.0001, one-way ANOVA; scale bars, 50 μm.

### Extended Data Fig. 4

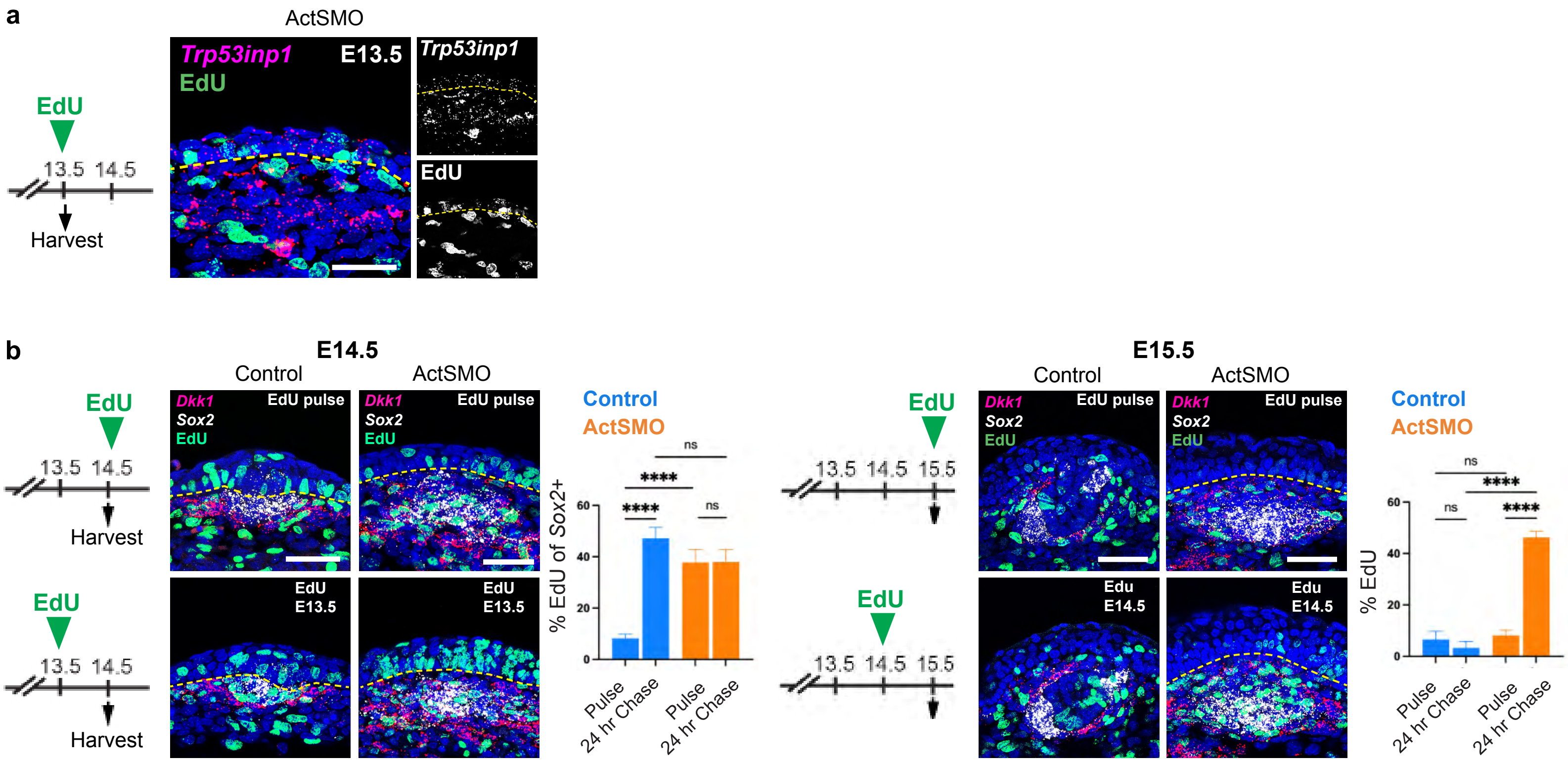


**Extended Data Fig. 4. DC cells are immediate progeny of a cell division in both wildtype and ActSMO embryos. a,** FISH stained skin of E13.5 ActSMO mutant after EdU was given for 1.5 hours prior to harvest (n=3). **b,** Left, FISH stained sections from either E14.5 (left) or E15.5 (right) control and ActSMO mutants that were either pulsed with EdU for 1.5 hours or 24 hours prior to harvest. Bar graphs show quantification of % EdU+ cells in *Sox2*+ cells for pulse and pulse-chase experiments in control or ActSMO conditions (n=4 per EdU regimen per time point). Data as mean± SEM; *P<0.05, **P<0.01, ***P<0.001, ****P<0.0001, one-way ANOVA; ns, not significant; scale bars, 50 μm.

### Extended Data Fig. 5

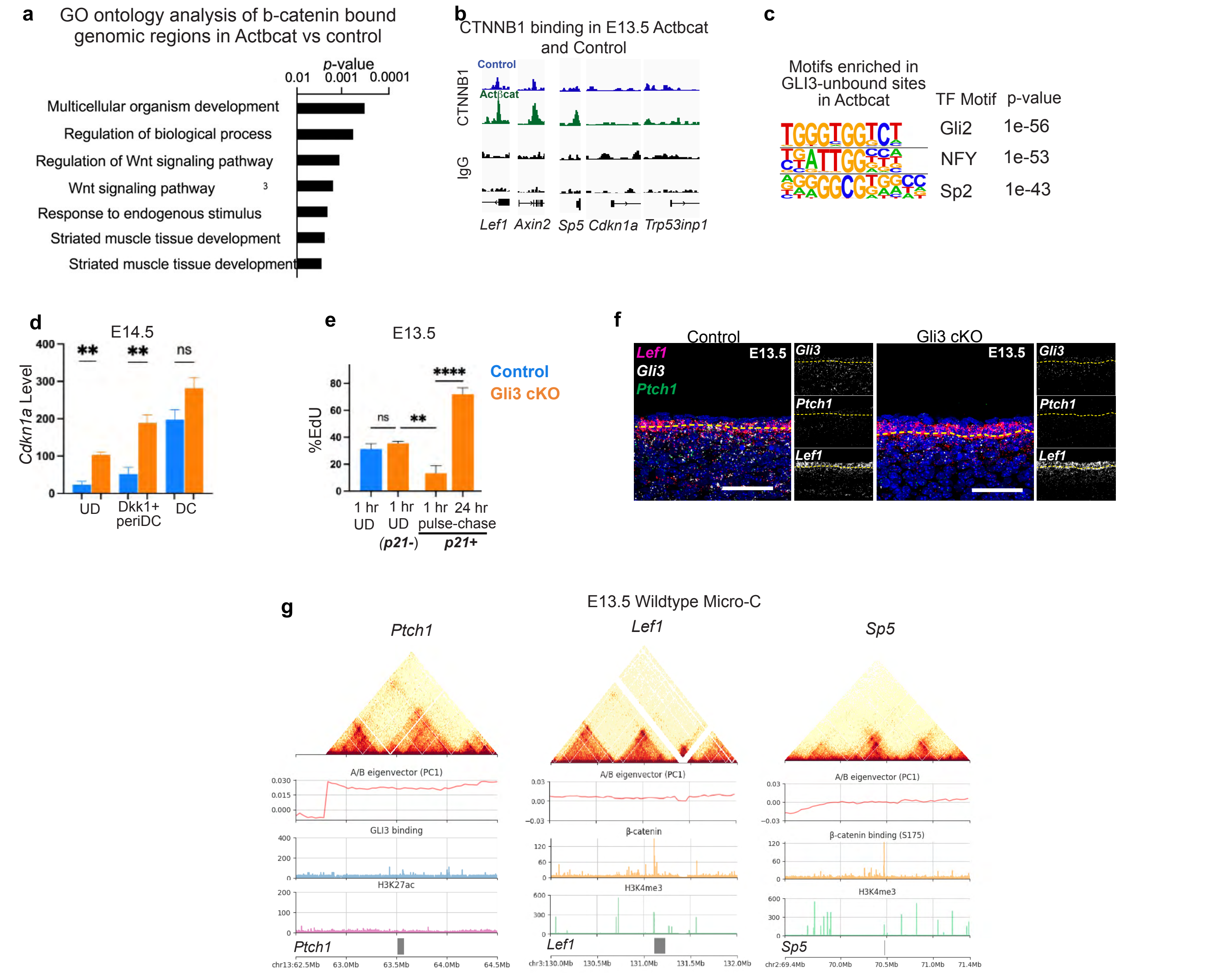


**Extended Data Fig. 5. GLI3 loss results in early cell-cycle exit in DC progenitors in the absence of SHH but does not repress many SHH target genes. a,** GO analysis of β–catenin–bound regions in Actβcat vs. control shows enrichment of Wnt signaling pathways but not cell cycle exit related pathways (p = 0.001 to 0.0001). **b**, CUT&RUN genome browser tracks of CTNNB1-bound regions in E13.5 Actβcat and control dermal cells at indicated genes, including Wnt targets (e.g., *Axin2*, *Lef1*, *Sp5*) and cell-cycle exit genes. **c,** Motif enrichment of GLI3-unbound sites in Actβcat skin, showing correspondence with GLI sites. **d,** FISH quantification of *Cdkn1a* transcript levels in E14.5 control vs Gli3 cKO dermis. **e,** %EdU of UD cells in E13.5 control or Gli3 cKO embryos pulsed with EdU for either 1 hour or 24 hours (E12.5) prior to harvest and parsed by *Cdkn1a* expression status. **f**, FISH skin sections stained for *Gli3*, *Ptch1*, and *Lef1* in control and Gli3 cKO embryos. **g**, E13.5 wildtype Micro-C contact maps with A/B eigenvector, GLI3 (leftmost) or CTNNB1 (middle, right) binding and H3K4me3 binding at indicated loci (*Ptch1*, *Lef1, Sp5* loci. Data as mean± SEM; *P<0.05, **P<0.01, ***P<0.001, ****P<0.0001, one-way ANOVA; scale bars, 50 µm.

**Extended Data Fig. 6**

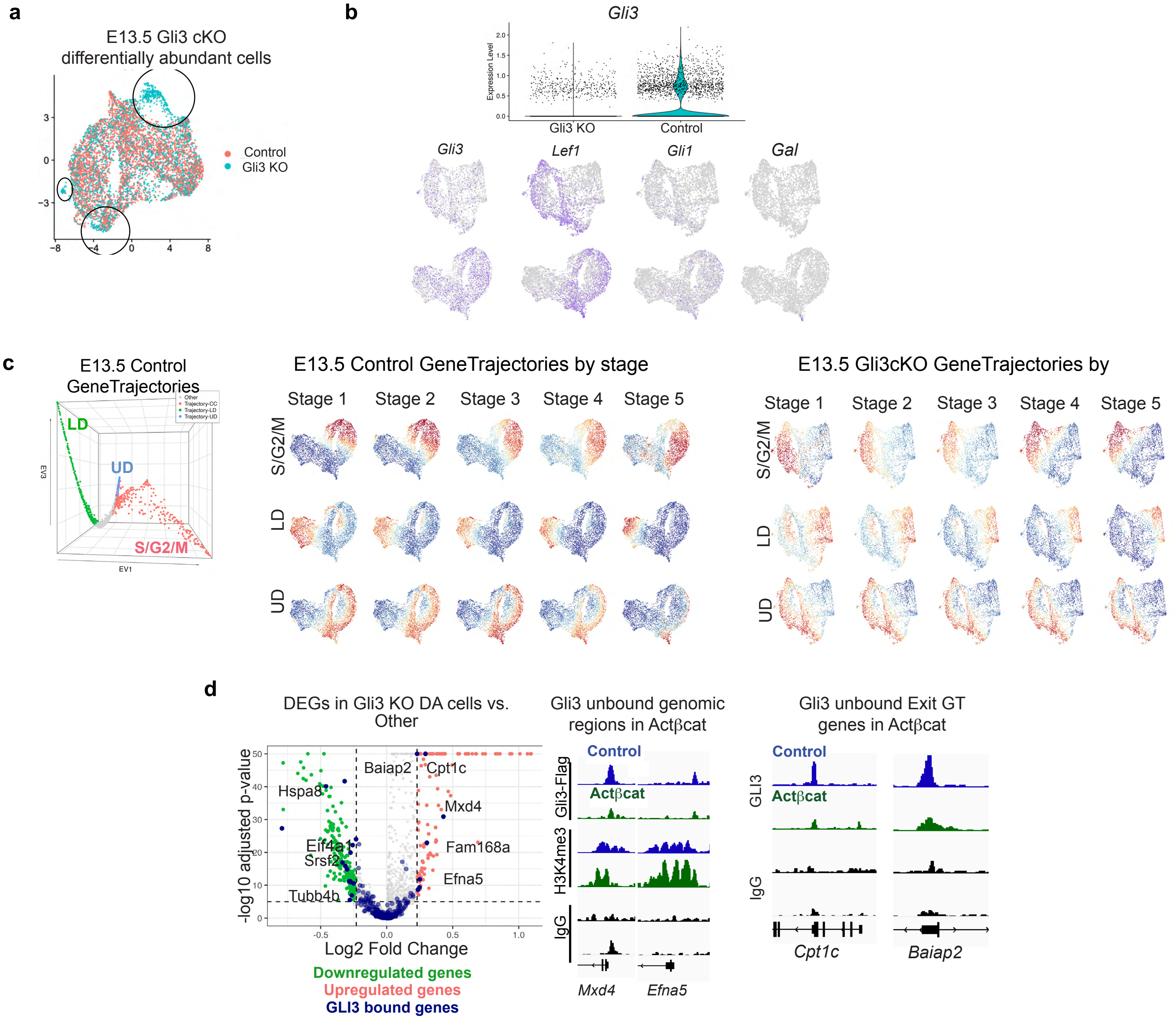


**Extended Data Fig. 6. Loss of dermal Gli3 results in an Exit gene trajectory without DC genes. a,** UMAPs of E13.5 control and Gli3 cKO dermis colored by condition (n=1). **b,** Violin plot of *Gli3* expression in control and Gli3 cKO dermal scRNA-seq data and dermal UMAPs colored by expression of indicated genes. **c,** Diffusion map of E13.5 control GTs with control UMAPs colored by indicated control GT stages (left) or Gli3 cKO UMAPs colored by mutant GT stages (right). **d,** Volcano plot of differentially expressed genes between differentially abundant cells in the Gli3 cKO vs control dermis with genes that overlap with GLI3-bound genes in the Actbcat mutant colored in navy blue (n=3). Right, CUT&RUN genome browser tracks showing GLI3 binding in E13.5 Actβcat and control at *Mxd4* and *Efna5* and Exit GT genes, *Cpt1c* and *Baibp2* (n=3).

### Extended Data Fig. 7

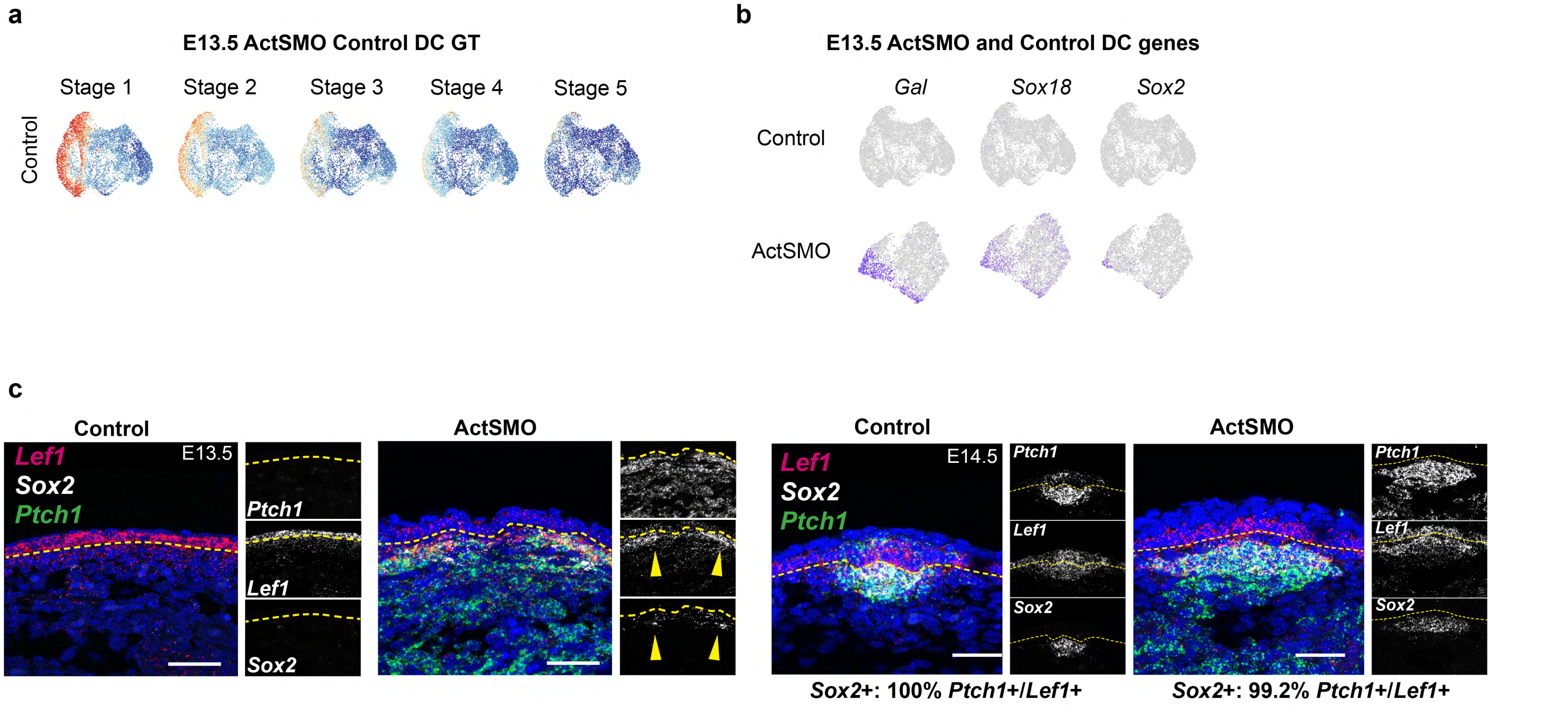


**Extended Data Fig. 7. SHH induces DC gene expression in Wnt-active cells throughout the upper dermis. a,** E13.5 ActSMO DC GT stages projected onto E13.5 paired control dermal scRNA-seq UMAPs showing a lack of most DC GT stages. **b,** E13.5 control and ActSMO dermal UMAPs colored by indicated DC marker genes. **c,** FISH images of E13.5 and E14.5 control and ActSMO mutant skin stained for *Ptch1, Lef1* and *Sox2* showing that virtually all *Sox2*+ cells express both *Ptch1* and *Lef1* while cells expressing *Ptch1* or *Lef1* alone do not express Sox2 (n=5). Scale bars, 50 µm.

**Extended Data Fig. 8**

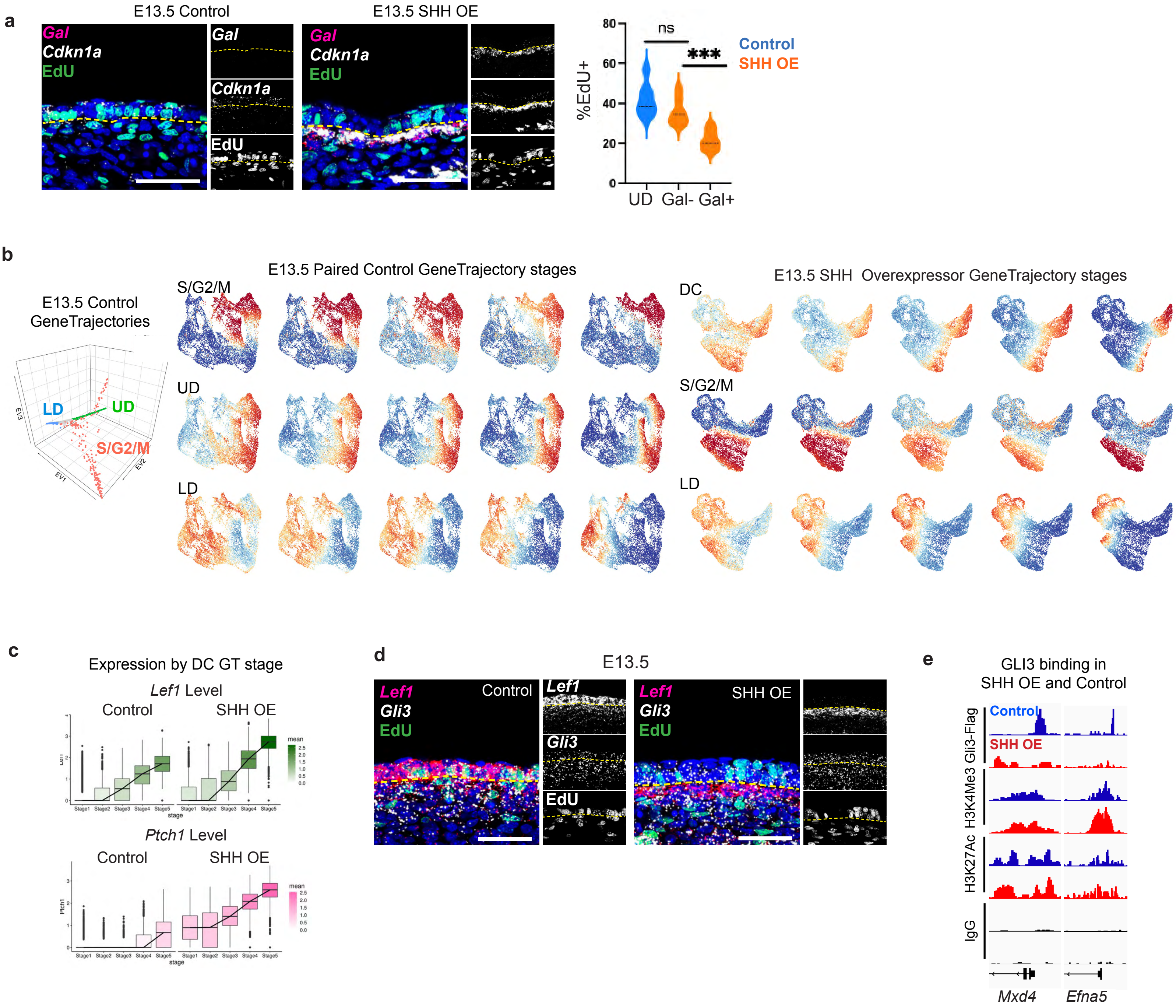


**Extended Data Fig. 8. Covarying SHH and Wnt gradients is sufficient to synchronize cell cycle exit with DC genes. a,** FISH stained images of E13.5 control or SHH OE skin showing colocalization of *Cdkn1a* with DC marker, *Gal* (n=3). Right, quantification of %EdU in E13.5 control UD or SHH OE UD parsed by *Gal* expresion (n=2). Note, *Gal* is not expressed in control at E13.5. **b,** Diffusion map of control GTs; UMAPs colored by indicated GT stages of E13.5 paired control (left) or GT stages for SHH OE (right). **c,** Quantification of *Lef1* or *Ptch1* expression across DC GT stages for E13.5 control and SHH OE mutant dermal cells. **d,** FISH stained sections showing *Gli3*, *Lef1*, and EdU in E13.5 control and SHH OE skin (n=3). **e,** CUT&DRUN track from E13.5 control or SHH OE skin showing binding by the indicated antibodies at *Mxd4* or *Efna5* genes (n=2). Data as mean± SEM; *P<0.05, **P<0.01, ***P<0.001, ****P<0.0001, one-way ANOVA; ns, not significant; scale bars, 50 µm.

**Extended Data Fig. 9**

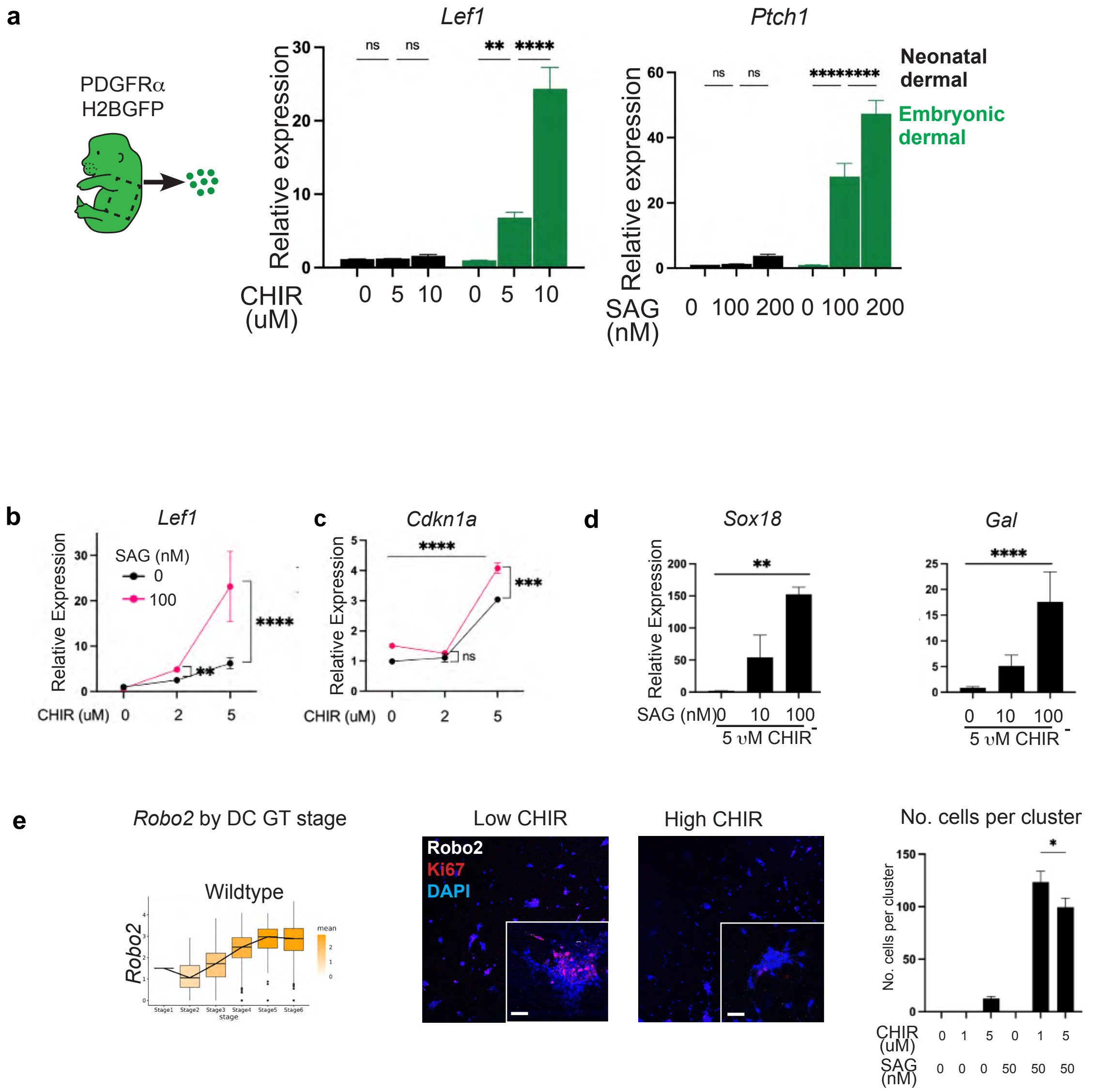


**Extended Data Fig. 9. Modulating levels of Wnt and SHH activity in vitro recapitulates in vivo molecular dynamics and DC changes. a,** qPCR of *Lef1* or *Ptch1* gene expression in response to varying doses of CHIR or SAG treatment for 48 hours using either neonatal (P0-P1) or E13.5 embryonic PDGFRαH2BGFP+ dermal cells (n=6). **b,** qPCR for *Lef1* or **c,** *Cdkn1a* or **d,** DC genes from embryonic dermal cells cultured for 48 hours with CHIR and/or SAG (n=3). **e,** *Robo2* expression by DC GT stage from E14.5 wildtype scRNA-seq data showing highest levels at the terminal DC stage. 3D dermal cultures grown in a collagen matrix for 48 hours with either low (1 μM) or high (5 μM) CHIR and stained for Ki67 and Robo2 (n=3). Right, quantification of cell number per cluster under different CHIR and/or SAG conditions (n=3). Data as mean± SEM; *P<0.05, **P<0.01, ***P<0.001, ****P<0.0001, one-way ANOVA; ns, not significant; scale bars, 50 μm.
